# Dual-color GRAB sensors for monitoring spatiotemporal serotonin release *in vivo*

**DOI:** 10.1101/2023.05.27.542566

**Authors:** Fei Deng, Jinxia Wan, Guochuan Li, Hui Dong, Xiju Xia, Yipan Wang, Xuelin Li, Chaowei Zhuang, Yu Zheng, Laixin Liu, Yuqi Yan, Jiesi Feng, Yulin Zhao, Hao Xie, Yulong Li

## Abstract

The serotonergic system plays important roles in both physiological and pathological processes, and is a widely used therapeutic target for many psychiatric disorders. Although several genetically encoded GFP-based serotonin (5-HT) sensors were recently developed, their sensitivities and spectral profiles are relatively limited. To overcome these limitations, we optimized green fluorescent G-protein-coupled receptor (GPCR)-activation-based 5-HT (GRAB_5-HT_) sensors and developed a new red fluorescent GRAB_5-HT_ sensor. These sensors have excellent cell surface trafficking, high specificity, sensitivity, and spatiotemporal resolution, making them suitable for monitoring 5-HT dynamics *in vivo*. Besides recording subcortical 5-HT release in freely moving mice, we observed both uniform and gradient 5-HT release in the mouse dorsal cortex with mesoscopic imaging. Finally, we performed dual-color imaging and observed seizure-induced waves of 5-HT release throughout the cortex following calcium and endocannabinoid waves. In summary, these 5-HT sensors can offer valuable insights regarding the serotonergic system in both physiological and pathological states.

## Introduction

Serotonin (5-HT) is an important monoamine signaling molecule present virtually throughout the body, widely regulating neural activity and other key biological processes^1^. In the central nervous system (CNS), 5-HT is an intensively studied neurotransmitter involved in a wide range of neurobiological processes such as emotion, learning and memory, reward, appetite, and the sleep-wake cycle^1–3^. Moreover, impaired 5-HT transmission is associated with a broad range of CNS disorders, including anxiety, addiction, depression, and epilepsy^4–6^. As a consequence, many psychotropic and psychedelic drugs have been developed to act on the serotonergic system in the CNS^7^. The primary source of 5-HT in the CNS is serotonergic neurons in the brainstem, which innervate most of the regions throughout the brain to drive various functions; moreover, these neurons are highly heterogeneous with respect to their transcriptomics and projection patterns^8–12^. To encode these widespread 5-HT signals into specific downstream signaling pathways, 15 different 5-HT receptor (5-HTR) subtypes have evolved^1^, with half-maximal effective concentration (EC_50_) values ranging from nanomolar to micromolar^1^^3^. Understanding the serotonergic system in both physiological and pathological processes requires the ability to directly monitor 5-HT dynamics in behaving animals in real time, which in turn requires highly sensitive detection tools. However, given the anatomical and functional complexities of the serotonergic system, classic detection methods such as microdialysis and fast-scan cyclic voltammetry (FSCV) lack the simultaneous high spatiotemporal resolution, specificity, sensitivity, and minimal invasiveness needed for the *in vivo* detection of 5-HT^14–16^.

Recent advances in genetically encoded fluorescent 5-HT sensors have led to overall optimal tools that surpass classic methods^17–20^; however, these sensors have not yet hit the proverbial “sweet spot” with respect to balancing apparent affinity with the magnitude of the response. Specifically, sensors based on GPCRs, including GRAB_5-HT1.0_ (ref.^17^), PsychLight2 (ref.^18^), and sDarken (ref.^19^), have high affinity for 5-HT but produce only a modest change in fluorescence. On the other hand, the periplasmic binding protein (PBP)-based sensor iSeroSnFR (ref.^20^) has a relatively large response, but low affinity to 5-HT. Thus, monitoring 5-HT dynamics *in vivo* requires a more sensitive, high-affinity sensor that produces a sufficiently large response.

In the brain, the serotonergic system interacts with other neurotransmitters and neuromodulators^21^; thus, simultaneously imaging 5-HT and other neurochemicals can provide valuable information regarding the regulation of cognitive functions controlled by these signaling processes. Unfortunately, most existing sensors for neurochemicals contain a green fluorescent protein (GFP) as the fluorescent module, as do all genetically encoded 5-HT sensors, precluding combined imaging due to spectral overlap. Although a near-infrared 5-HT nanosensor based on single-wall carbon nanotubes has been reported^22^, it may not be suitable for use in living animals due to limited sensitivity. On the other hand, red-shifted sensors, such as a previously reported red calcium sensor^23, 24^, are compatible with other green fluorescent sensors and blue light-excitable actuators, with intrinsically superior optical properties—including deeper tissue penetration, reduced autofluorescence, and low phototoxicity—due to their longer excitation wavelengths. Thus, red-shifted 5-HT sensors suitable for *in vivo* imaging, particularly multiplexed imaging, are urgently needed.

Here, we report a series of green and red fluorescent 5-HT sensors generated by transplanting the third intracellular loop (ICL3)—containing circular permutated enhanced GFP (cpEGFP) or cpmApple—from existing green and red GRAB sensors into the 5-HTR4 subtype. The green fluorescent sensor, gGRAB_5-HT3.0_ (g5-HT3.0), produces a ∼1,300% increase in fluorescence in response to 5-HT, making it vastly superior to existing green fluorescent 5-HT sensors both *in vitro* and *in vivo*. The red-shifted sensor, rGRAB_5-HT1.0_ (r5-HT1.0), produces a >300% increase in fluorescence in response to 5-HT and is also suitable for both *in vitro* and *in vivo* applications. Using mesoscopic imaging in mice expressing g5-HT3.0, we found that 5-HT is released in a gradient along the anterior-to-posterior axis in the mouse dorsal cortex upon optogenetic stimulation of serotonergic neurons in the dorsal raphe nucleus (DRN), but is released with spatial homogeneity during the sleep-wake cycle. Finally, dual-color mesoscopic imaging revealed cortex-wide 5-HT waves that followed calcium and endocannabinoid (eCB) waves in a mouse seizure model. Thus, these improved dual-color GRAB_5-HT_ sensors are powerful tools for monitoring serotonin release *in vivo*, providing valuable new insights into the serotonergic system.

## Results

### Development and optimization of new GRAB_5-HT_ sensors

To expand the dynamic range and spectral profile of GRAB_5-HT_ sensors, we first systematically searched for the most suitable GPCR scaffold by transplanting the ICL3 from existing green and red fluorescent GRAB sensors into a wide range of 5-HT receptor subtypes (Fig. 1a). For green fluorescent sensors, we used green GRAB_5-HT1.0_ (g5-HT1.0)^17^ or GRAB_NE1m_ (ref.^25^) as the ICL3 donor (including cpEGFP and surrounding linker sequences). Possibly due to the conserved structures among the class A family of GPCRs, we successfully obtained new candidate sensors that yielded a >50% increase in fluorescence (ΔF/F_0_) in response to 10 µM 5-HT following the screening of replacement sites. One green fluorescent candidate based on 5-HTR4 had a higher response than the original g5-HT1.0 sensor and was named g5-HT1.1. To develop red fluorescent 5-HT sensors, we transplanted the ICL3 (including cpmApple and the linker sequences) from rGRAB_DA1m_ (ref.^26^) into the 5-HTR2C, 5-HTR4, and 5-HTR6 scaffolds—we chose these three subtypes based on their high performance when developing the green fluorescent 5-HT sensors. Once again, the top candidate was based on the 5-HTR4 subtype, with a ∼40% ΔF/F_0_; we named this sensor r5-HT0.1 (Fig. 1a).

**Fig. 1.**
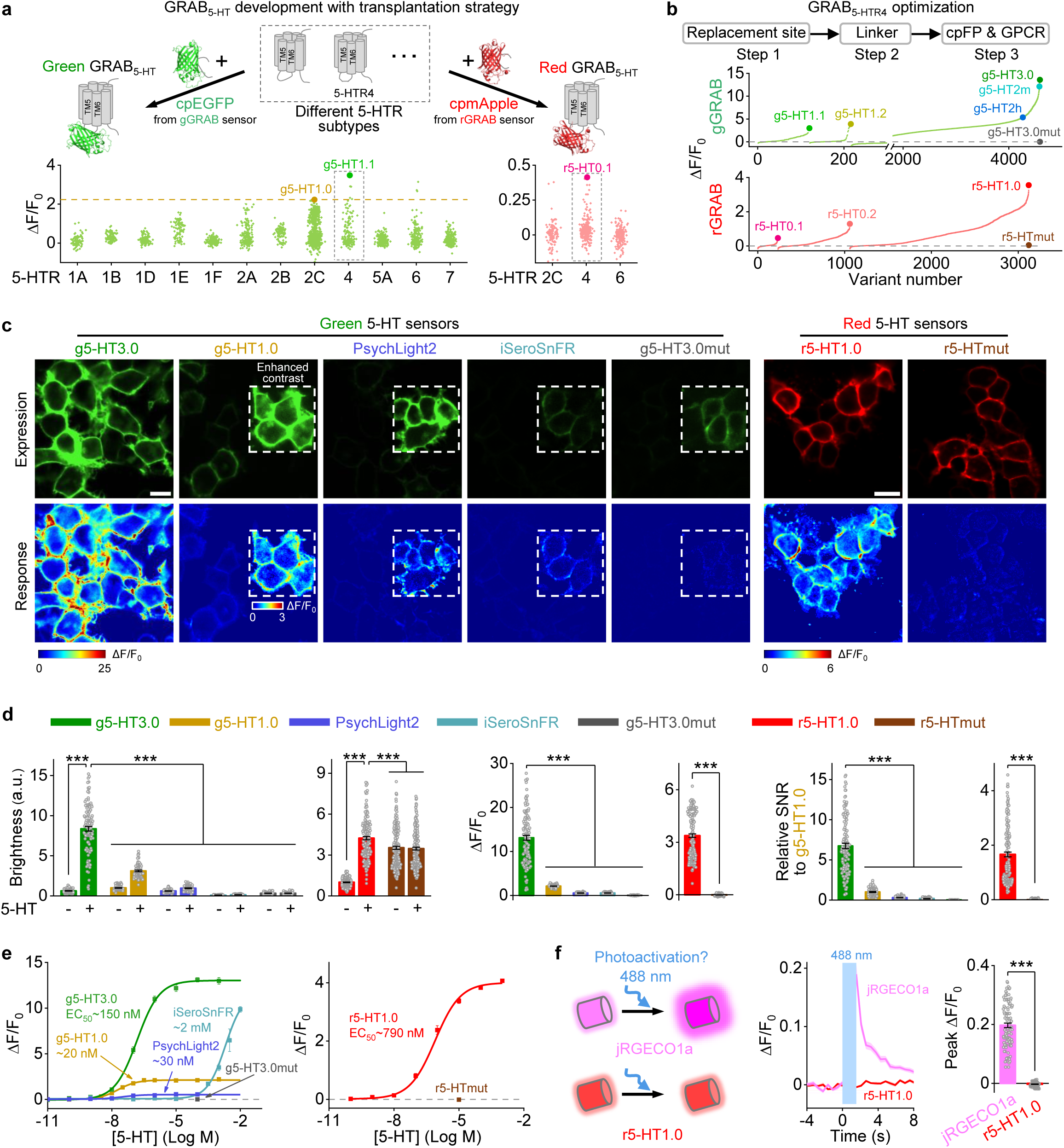
| Development of improved green fluorescent 5-HT sensors and new red 5-HT sensors. **a**, Schematic illustrating the strategy for developing new GRAB_5-HT_ sensors (top). Performance of sensor candidates based on different receptor subtypes for green (bottom left) and red 5-HT sensors (bottom right). The dashed horizontal line represents g5-HT1.0 response (bottom left), and the best candidates in green and red sensors are denoted by enlarged green and pink dots, respectively. **b**, Optimization of the replacement site, linker, cpFP and GPCR. Responses to 10 μM 5-HT of various candidates are presented, and different versions are indicated with enlarged dots. **c**, Representative images of sensors’ expression (top, with 5-HT) and response (bottom) to 5-HT in HEK293T cells. Insets with white dashed outlines in images have either enhanced contrast (top) or different pseudocolor scales (bottom). 100 μM 5-HT for green sensors and 10 μM 5-HT for red sensors. Scale bar, 20 μm. **d**, Group summary of the brightness (left), peak ΔF/F_0_ (middle) and SNR (right) of different 5-HT sensors. The SNR of all sensors were relative to g5-HT1.0; a.u., arbitrary units, the basal brightness of g5-HT1.0 is set as 1. *n* = 119 cells from 3 coverslips (hereafter denoted as 119/3) for g5-HT3.0, 82/3 for g5-HT1.0, 64/3 for PsychLight2, 139/3 for iSeroSnFR, 92/3 for g5-H3.0mut, 159/5 for r5-HT1.0 and 191/5 for r5-HTmut; 100 μM 5-HT for green sensors and 10 μM 5-HT for red sensors. (One-way ANOVA followed by Tukey’s multiple-comparison tests for green sensors; for brightness, *F*_9,982_ = 600.2, *P* = 0, post hoc test: *P* <10^-8^ for g5-HT3.0 with 5-HT versus g5-HT3.0 without 5-HT and other sensors with or without 5-HT; for peak ΔF/F_0_, *F*_4,491_ = 387.1, *P* = 2.76×10^-150^, post hoc test: *P* <10^-8^ for g5-HT3.0 versus other sensors; for relative SNR, *F*_4,491_ = 285.7, *P* = 1.13×10^-1^^26^, post hoc test: *P* <10^-6^ for g5-HT3.0 versus other sensors. One-way ANOVA followed by Tukey’s multiple-comparison tests for brightness of red sensors, *F*_3,696_ = 178.3, *P* = 9.26×10^-86^, post hoc test: *P* <10^-5^ for r5-HT1.0 with 5-HT versus r5-HT1.0 without 5-HT and r5-HTmut with or without 5-HT. Two-tailed Student’s *t*-test for r5-HT1.0 and r5-HTmut; for peak ΔF/F_0_, *P* = 3.13×10^-72^; for relative SNR, *P* = 2.67×10^-43^.) **e**, Dose-response curves of different 5-HT sensors. *n* = 3 wells for each sensor with 300–500 cells per well. **f**, Schematic illustrates the photoactivation properties of jRGECO1a and r5-HT1.0 (left), representative traces (middle) and group summary of peak ΔF/F_0_ (right) in response to blue light (488 nm, without imaging) in cells expressing jRGECO1a or r5-HT1.0. *n* = 105/4 for jRGECO1a and 88/4 for r5-HT1.0. (Two-tailed Student’s *t*-test, *P* = 2.07×10^-48^). Data are shown as mean ± SEM in **d–f**, with the error bars or shaded regions indicating the SEM, \*\*\**P* < 0.001.

To improve the sensors’ sensitivity, then we optimized g5-HT1.1 and r5-HT0.1 by performing saturation mutagenesis of critical residues believed to affect structural coupling, fluorescence intensity^27^, protein folding^28^, and the 5-HT induced conformational change^29^. Iterative optimization by screening more than 4,500 candidates yielded two intermediate green fluorescent sensors called g5-HT2h and g5-HT2m, followed by the final sensor g5-HT3.0 (Fig. 1b,c and Extended Data Figs. 1a–c). When expressed in HEK293T cells, these green fluorescent 5-HT sensors showed good trafficking to the cell surface and a robust increase in fluorescence in response to 5-HT application (Fig. 1c,d and Extended Data Fig. 2a–c). In response to 100 µM 5-HT, the change in fluorescence of g5-HT3.0 (ΔF/F_0_: ∼1,300%) was 6–20-fold larger than the response measured for previously developed 5-HT sensors, g5-HT1.0, PsychLight2, and iSeroSnFR^17, 18, 20^. Moreover, the g5-HT3.0 sensor is significantly brighter, with a higher signal-to-noise ratio (SNR) than the other green fluorescent sensors (Fig. 1d). Dose-response curves showed that g5-HT3.0 produces a larger ΔF/F_0_ (i.e., is more sensitive to 5-HT) than previous sensors over a wide range of concentrations, with an EC_50_ value of ∼150 nM (Fig. 1e). We also generated a 5-HT insensitive version of the g5-HT3.0 sensor, named g5-HT3.0mut, by introducing the D131^3.32^F mutation^30, 31;^ we confirmed that g5-HT3.0mut localizes to the plasma membrane but does not respond to 5-HT even at 100 µM (Fig. 1b–e and Extended Data Fig. 1a,c).

Similarly, we generated a red fluorescent 5-HT sensor called r5-HT1.0 by screening >3,000 candidates, and we generated a 5-HT insensitive version, named r5-HTmut, by introducing the D131^3.32^Q and D149^3.50^H mutations (Fig. 1b–e and Extended Data Fig. 1d,f). Both r5-HT1.0 and r5-HTmut localized to the plasma membrane when expressed in cultured HEK293T cells (Fig. 1c). Application of 10 µM 5-HT to cells expressing r5-HT1.0 elicited a ∼330% increase in fluorescence, but had no effect on cells expressing r5-HTmut (Fig. 1c,d). The EC_50_ of r5-HT1.0 was ∼790 nM (Fig. 1e). Moreover, unlike the cpmApple-based calcium sensor jRGECO1a—in which blue light causes an increase in fluorescence^24, 32^—we found that blue light had no detectable effect on r5-HT1.0 (Fig. 1f).

### Characterization of GRAB_5-HT_ sensors in cultured cells

Next, we characterized the pharmacology, specificity, spectra, and kinetics of our new 5-HT sensors expressed in HEK293T cells. We found that both g5-HT3.0 and r5-HT1.0 inherited the pharmacological specificity of the parent 5-HTR4 receptor, as their 5-HT induced responses were blocked by the 5-HTR4-specific antagonist RS 23597-190 (RS), but not the 5-HTR2C-specific antagonist SB 242084 (SB); in addition, both sensors were unaffected by application of a wide range of signaling molecules, including the 5-HT precursor, 5-HT metabolite, and a variety of other neurotransmitters and neuromodulators (Fig. 2a and Extended Data Fig. 3a,b). The emission spectra of the green fluorescent (with a peak at 520 nm) and red fluorescent (with a peak at 595 nm) sensors are well-separated, and we measured 1-photon/2-photon excitation peaks at 505/920 nm for g5-HT3.0 and 560/1050 nm for r5-HT1.0 (Fig. 2b and Extended Data Fig. 4a,b). We also measured a 425-nm isosbestic point for g5-HT3.0 under 1-photon excitation. With respect to the sensors’ kinetics, we measured the on rate (τ_on_) by locally puffing 10 μM 5-HT on the cells and the off rate (τ_off_) by puffing the 5-HTR4 antagonist RS in the continued presence of 10 μM 5-HT (Fig. 2c,d), revealing sub-second τ_on_ rates and faster τ_off_ rates than our previously reported g5-HT1.0 sensor^17^, with mean τ_off_ rates of 1.66 s, 1.90 s, 0.38 s, and 0.51 s for g5-HT3.0, g5-HT2h, g5-HT2m, and r5-HT1.0, respectively (Fig. 2e and Extended Data Fig. 4c).

**Fig. 2.**
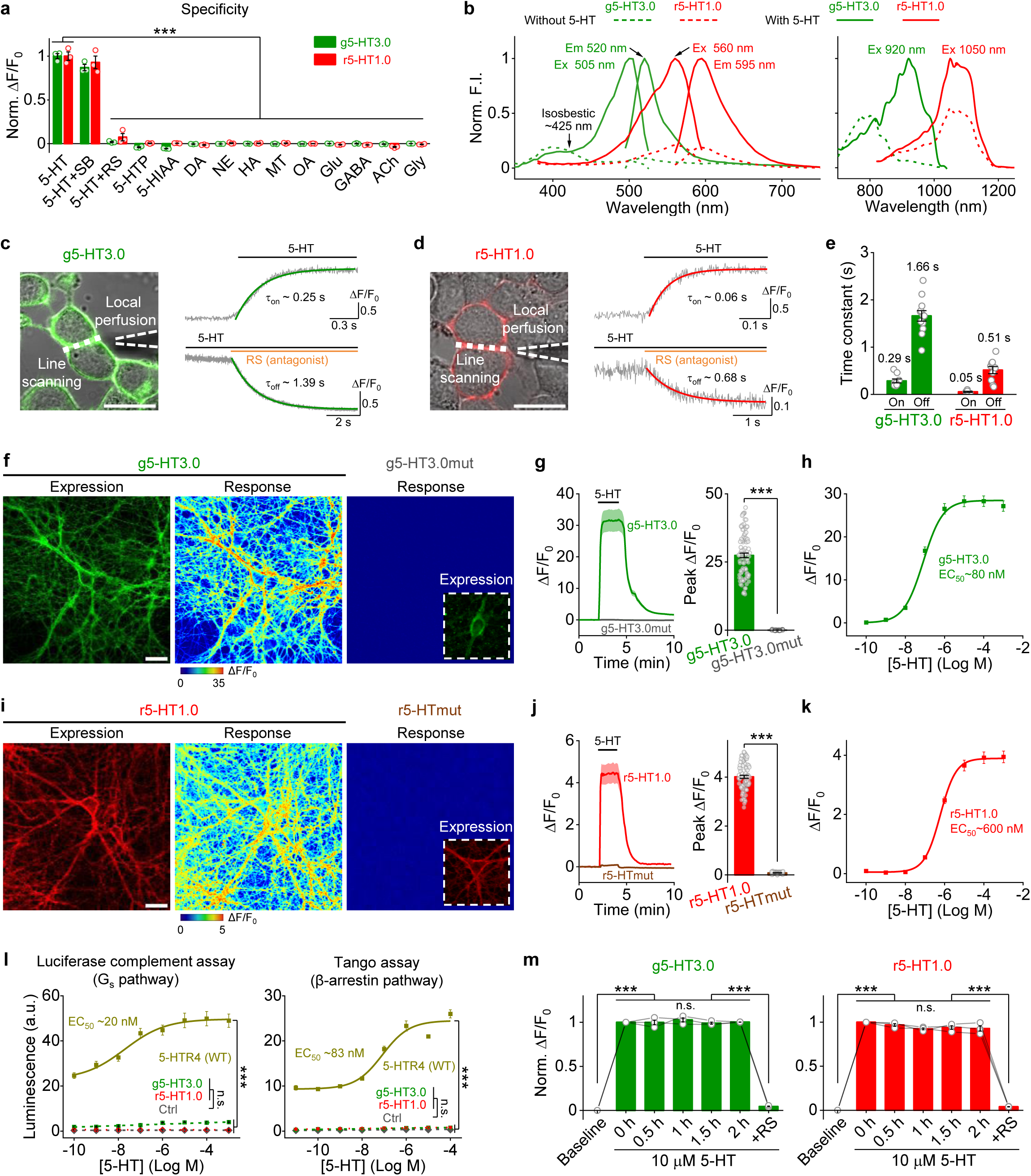
| Characterization of 5-HT sensors in HEK293T cells and cultured rat cortical neurons. **a**, Normalized ΔF/F_0_ of g5-HT3.0 and r5-HT1.0 in response to different compounds (each at 10 μM except RS at 100 μM). 5-HTP, 5-hydroxytryptophan; 5-HIAA, 5-hydroxyindole acetic acid; DA, dopamine; NE, norepinephrine; HA, histamine; MT, melatonin; OA, octopamine; Glu, glutamate; GABA, gamma-aminobutyric acid; ACh, acetylcholine; Gly, glycine. Norm., normalized. *n* = 3 wells per group, 200–500 cells per well. (For g5-HT3.0, *F*_13,28_ = 745.7, *P* = 5.74×10^-32^, post hoc test: *P* = 0 for 5-HT versus 5-HT and RS, and other compounds; for r5-HT1.0, *F*_13,28_ = 180.6, *P* = 2.02×10^-23^, post hoc test: *P* = 0 for 5-HT versus 5-HT and RS, and other compounds.) **b**, One-photon excitation (Ex) and emission (Em) spectra and two-photon excitation spectra of g5-HT3.0 and r5-HT1.0 in the absence (dashed line) and presence of 10 μM 5-HT (solid line). F.I., fluorescence intensity. **c–e**, Kinetic of g5-HT3.0 and r5-HT1.0 in cultured HEK293T cells. Illustration of the local puffing system (**c**,**d**, left). Representative traces of sensor fluorescence increase to 5-HT puffing (**c**,**d**, top right) and decrease to RS puffing (**c**,**d**, bottom right). Group summary of on and off kinetics (**e**). *n* = 10 cells from 3 coverslips (short for 10/3) for g5-HT3.0 on kinetics, 12/4 for g5-HT3.0 off kinetics, 9/3 for r5-HT1.0 on kinetics, 12/4 for r5-HT1.0 off kinetics. **f**, Representative images showing the expression and responses of g5-HT3.0 and g5-HT3.0mut to 100 μM 5-HT in cultured rat cortical neurons. The inset in the g5-HT3.0mut response image shows the contrast-enhanced expression image. **g**, Representative traces and peak response summary of g5-HT3.0 and g5-HT3.0mut in response to 100 μM 5-HT. *n* = 96 ROIs from 5 coverslips (short for 96/5) for g5-HT3.0 and 92/5 for g5-HT3.0mut. (*P* = 1.40×10^-53^ for g5-HT3.0 versus g5-HT3.0mut.) **h**, The dose-response curve of g5-HT3.0. *n* = 76/4. **i**, Representative images showing the expression and responses of r5-HT1.0 and r5-HTmut to 10 μM 5-HT. **j**, Representative traces and peak response summary of r5-HT1.0 and r5-HTmut in response to 10 μM 5-HT. *n* = 80/4 for r5-HT1.0 and 60/3 for r5-HTmut. (*P* = 4.46×10^-70^ for r5-HT1.0 versus r5-HTmut.) **k**, The dose-response curve of r5-HT1.0. *n* = 80/4. **l**, Downstream coupling tests. WT, wild type; Ctrl, control, without expression of wild type 5-HTR4 or sensors; a.u., arbitrary units. *n* = 3 wells per group, 200–500 cells per well. (For luciferase complement assay, *F*_3,8_ = 256, *P* = 2.77×10^-8^, post hoc test: *P* = 0 and 0.37 for g5-HT3.0 versus 5-HTR4 (WT) and Ctrl in 1 mM 5-HT, respectively, *P* = 0 and 1 for r5-HT1.0 versus 5-HTR4 (WT) and Ctrl, respectively; for Tango assay, *F*_3,8_ = 766.4, *P* = 3.55×10^-10^, post hoc test: *P* = 0 and 0.89 for g5-HT3.0 versus 5-HTR4 (WT) and Ctrl in 100 μM 5-HT, respectively, *P* = 0 and 0.86 for r5-HT1.0 versus 5-HTR4 (WT) and Ctrl, respectively.) **m**, Normalized ΔF/F_0_ of g5-HT3.0 and r5-HT1.0 in response to the 2-h application of 10 μM 5-HT, followed by 100 μM RS. *n* = 3 wells for each sensor. (For g5-HT3.0, *F* = 359.8, *P* = 0.034, post hoc test: *P* = 1.29×10^-6^ for baseline versus 0 h, *P* = 1.76×10^-6^ for 2.0 h versus RS, *P* = 1, 0.77, 1, 1 for 0 h versus 0.5 h, 1 h, 1.5 h or 2.0 h, respectively; for r5-HT1.0, *F* = 250.9, *P* = 0.04, post hoc test: *P* = 2.85×10^-6^ for baseline versus 0 h, *P* = 5.82×10^-6^ for 2.0 h versus RS, *P* = 0.95, 0.44, 0.66, 0.64 for 0 h versus 0.5 h, 1 h, 1.5 h or 2.0 h, respectively.) **a–e** and **l** tested in HEK293T cells; **f–k** and **m** tested in cultured rat cortical neurons. All scale bar, 20 μm. Data are shown as mean ± SEM in **a**,**e**,**g**,**h**,**j–m**, with the error bars or shaded regions indicating the SEM. One-way ANOVA (in **a**,**l**) and one-way repeated measures ANOVA (in **m**) followed by Tukey’s multiple-comparison tests; two-tailed Student’s *t*-test in **g**,**j**; \*\*\**P* < 0.001, n.s., not significant.

We then expressed the g5-HT3.0 and r5-HT1.0 sensors in cultured rat cortical neurons and found that they trafficked well to the cell membrane and were distributed throughout the soma, dendrites, and axon. Application of a saturating concentration of 5-HT induced a fluorescence increase of ∼2,700% for g5-HT3.0 and ∼400% for r5-HT1.0, but had no effect on neurons expressing g5-HT3.0mut or r5-HTmut (Fig. 2f,g,i,j). We also confirmed the sensors’ high specificity for 5-HT when expressed in cultured cortical neurons (Extended Data Fig. 3c–f), and we measured EC_50_ values of 80 nM, 70 nM, 2.4 µM, and 600 nM for g5-HT3.0, g5-HT2h, g5-HT2m, and r5-HT1.0, respectively (Fig. 2h,k and Extended Data Fig. 4d). Compared to previously reported GFP-based 5-HT sensors^17, 18, 20^, g5-HT3.0 is brighter and has a greater fluorescence change and a higher SNR when expressed in cultured neurons (Extended Data Fig. 5).

The steric hindrance of the bulky cpFP in sensors is likely to disturb the downstream coupling of GPCR^17, 25, 33^. To confirm that our 5-HT sensors do not couple to downstream signaling pathways—and therefore do not likely affect cell activity—we used the luciferase complement assay^34^ and the Tango assay^35, 36^ to measure the GPCR-mediated G_s_ and β-arrestin pathways, respectively. We found that g5-HT3.0, g5-HT2h, g5-HT2m, and r5-HT1.0 had negligible downstream coupling; in contrast, the wild-type 5-HTR4 receptor had high basal activity and robust, dose-dependent coupling (Fig. 2l and Extended Data Fig. 4e,f). In addition, our sensors do not undergo β-arrestin-mediated internalization or desensitization when expressed in cultured neurons, as the 5-HT elicited increase in fluorescence was stable for up to 2 hours in the continuous presence of 10 µM 5-HT (Fig. 2m and Extended Data Fig. 4g).

### Measuring endogenous 5-HT release in freely moving mice

To determine whether our newly developed red fluorescent sensor is suitable for *in vivo* imaging in freely behaving mice, we expressed either r5-HT1.0 or r5-HTmut in the basal forebrain (BF), which receives extensive DRN serotonergic projections^37^, and expressed the light-activated channel ChR2 (ref^.38, 39^) in serotonergic neurons in the DRN of *Sert-Cre* mice^40^ (Fig. 3a). Optical stimulation of the DRN induced time-locked transient increases in r5-HT1.0 fluorescence, the amplitude of which increased progressively with increasing stimulation duration; moreover, the selective serotonin transporter blocker fluoxetine further increased the amplitude of the response and prolonged the response’s decay kinetics (Fig. 3b–e). As expected, no response was measured for the 5-HT insensitive r5-HTmut sensor (Fig. 3b–e).

**Fig. 3.**
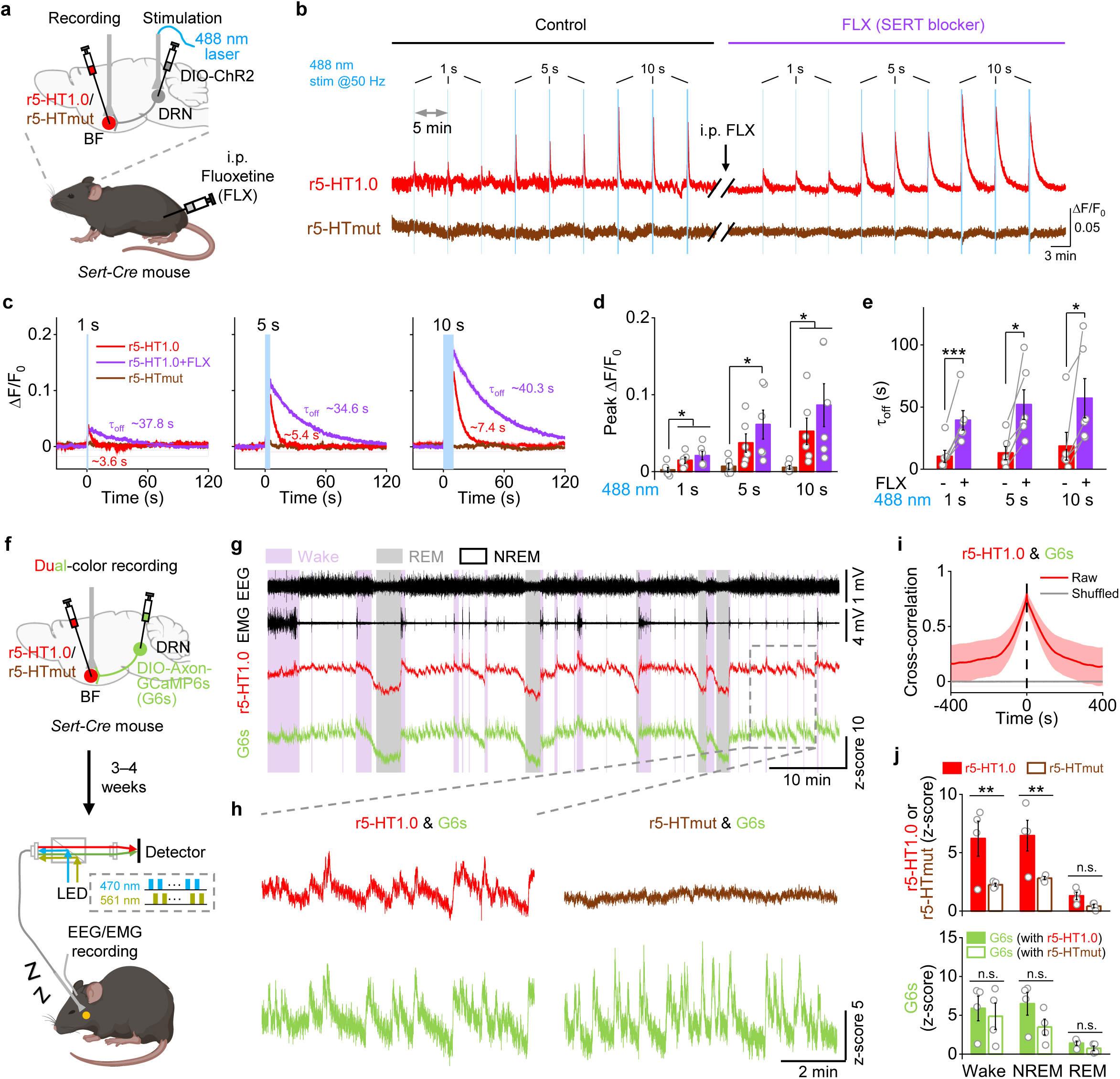
| Red GRAB_5-HT_ sensor can monitor endogenous 5-HT release in freely moving mice. **a**, Schematic depicting the fiber-photometry recording setup using red 5-HT sensors with optogenetic activation of DRN in *Sert-Cre* mice. **b**, Representative ΔF/F_0_ traces of r5-HT1.0 and r5-HTmut in response to optical stimulation in the DRN under different stimulation durations before or after fluoxetine (FLX) application. Blue shading, period for 488-nm stimulation. **c**, Averaged ΔF/F_0_ traces of r5-HT1.0 and r5-HTmut under different stimulation durations in an example mouse. **d**, Summarized peak responses of r5-HT1.0 and r5-HTmut under different stimulation durations. *n* = 6 mice for each treatment. (Two-tailed Student’s *t*-tests, for r5-HTmut versus r5-HT1.0, *P* = 0.030, 0.052, 0.041 under 1 s, 5 s, 10 s stimulation, respectively; for r5-HTmut versus r5-HT1.0+FLX, *P* = 0.016, 0.034, 0.033 under 1 s, 5 s, 10 s stimulation, respectively.) **e**, Summarized decay kinetics of r5-HT1.0 with or without FLX application under different stimulation durations. *n* = 6 mice for each treatment. (Paired *t*-tests for r5-HT1.0 and r5-HT1.0 + FLX, *P* = 4.44×10^-4^, 1.44×10^-2^, 3.19×10^-2^ for 1 s, 5 s and 10 s stimulation, respectively.) **f**, Schematic showing the setup for dual-color recording of r5-HT1.0 or r5-HTmut and GCaMP6s (G6s) during the sleep-wake cycle. **g**, Representative traces of simultaneous EEG, EMG, r5-HT1.0, and G6s recording during the sleep-wake cycle in freely behaving mice. Pink shading, wake state; gray shading, REM sleep. **h**, Zoom-in traces of r5-HT1.0 and G6s (from **g**) or r5-HTmut and G6s (mainly during the NREM sleep). **i**, The averaged cross-correlation between r5-HT1.0 and G6s signals during the sleep-wake cycle. **j**, Averaged responses of 5-HT sensors (red channel, r5-HT1.0 or r5-HTmut) and G6s (green channel). *n* = 4 mice for each group. (Two-way repeated measures ANOVA followed by Tukey’s multiple-comparison tests; for r5-HT1.0 versus r5-HTmut, post hoc test: *P* = 5.65×10^-3^, 9.22×10^-3^ and 0.47 during wake, NREM and REM sleep state, respectively; for G6s (with r5-HT1.0) versus G6s (with r5-HTmut), post hoc test: *P* = 0.56, 0.11 and 0.71 during wake, NREM and REM sleep state, respectively.) Data are shown as mean ± SEM in **c–e**,**i**,**j**, with the error bars or shaded regions indicating the SEM, \**P* < 0.05, \*\**P* < 0.01, \*\*\**P* < 0.001, n.s., not significant.

To test whether r5-HT1.0 is compatible with green fluorescent sensors, we expressed either r5-HT1.0 or r5-HTmut in the BF and expressed the axon-targeted green fluorescent calcium sensor axon-GCaMP6s (ref.^41^) in DRN serotonergic neurons, which project to the BF and regulate the sleep-wake cycle^17, 42^. We then performed dual-color fiber photometry recording in the BF while simultaneously recording the electroencephalography (EEG) and electromyography (EMG) signals in order to track the animal’s sleep-wake state (Fig. 3f). We found that both the r5-HT1.0 and GCaMP6s signals were higher during both the wake state and NREM (non-rapid eye movement) sleep than during REM sleep (Fig. 3g). In addition, r5-HT1.0 revealed oscillations in 5-HT levels (Fig. 3h). Moreover, we found that the r5-HT1.0 and GCaMP6s signals were temporally correlated, with no detectable lag, revealing the rapid kinetics of 5-HT release and high consistency between 5-HT release and the increase in presynaptic calcium (Fig. 3i). In contrast, the r5-HTmut signal was largely unchanged throughout the sleep-wake cycle and was significantly smaller than the r5-HT1.0 signal during the wake state and NREM sleep (Fig. 3j and Extended Data Fig. 6). Finally, we found no significant difference in the GCaMP6s signal between mice co-expressing r5-HTmut and mice co-expressing r5-HT1.0 (Fig. 3j).

To compare the performance of our optimized g5-HT3.0 sensor to previously reported 5-HT sensors, we also performed bilateral recordings in the BF during the sleep-wake cycle in mice expressing g5-HT3.0 in one hemisphere and g5-HT1.0, PsychLight2, or iSeroSnFR in the other hemisphere (Extended Data Fig. 7a,d,g). Consistent with our *in vitro* results, we found that the g5-HT3.0 sensor had significantly larger SNR during the wake state and NREM sleep compared to the other three sensors, as well as more robust oscillations measured during NREM sleep (Extended Data Fig. 7b,c,e,f,h,i).

### Mesoscopic imaging of 5-HT dynamics in the mouse dorsal cortex

5-HT also plays important roles in the cerebral cortex, for example regulating cognition and emotion in the prefrontal cortex^12, 37, 43^. Given that the cortex generally receives sparser serotonergic projections compared to subcortical regions such as the BF^44^, measuring 5-HT release in the cortex requires a highly sensitive 5-HT sensor. To imaging 5-HT dynamics throughout the whole dorsal cortex, we expressed the g5-HT3.0 sensor by injecting AAV in the transverse sinus^45^. We then measured g5-HT3.0 fluorescence throughout the cortex using mesoscopic imaging^46, 47^ in response to optogenetic stimulation of DRN serotonergic neurons expressing ChrimsonR^48^ (Fig. 4a). We found that light pulses induced transient increases in g5-HT3.0 fluorescence, with increasing stimulation frequency causing increasingly larger responses (Fig. 4b,c). As a negative control, no response was measured when we expressed a membrane-tethered EGFP (memEGFP) (Fig. 4 b,c and Extended Data Fig. 8g). Importantly, we found that treating mice with the SERT blocker fluoxetine caused a gradual increase in baseline 5-HT levels and slowed the decay rate of stimulation-induced transients; in contrast, the dopamine transporter (DAT) blocker GBR 12909 had no effect (Fig. 4d and Extended Data Fig. 8a–f).

**Fig. 4.**
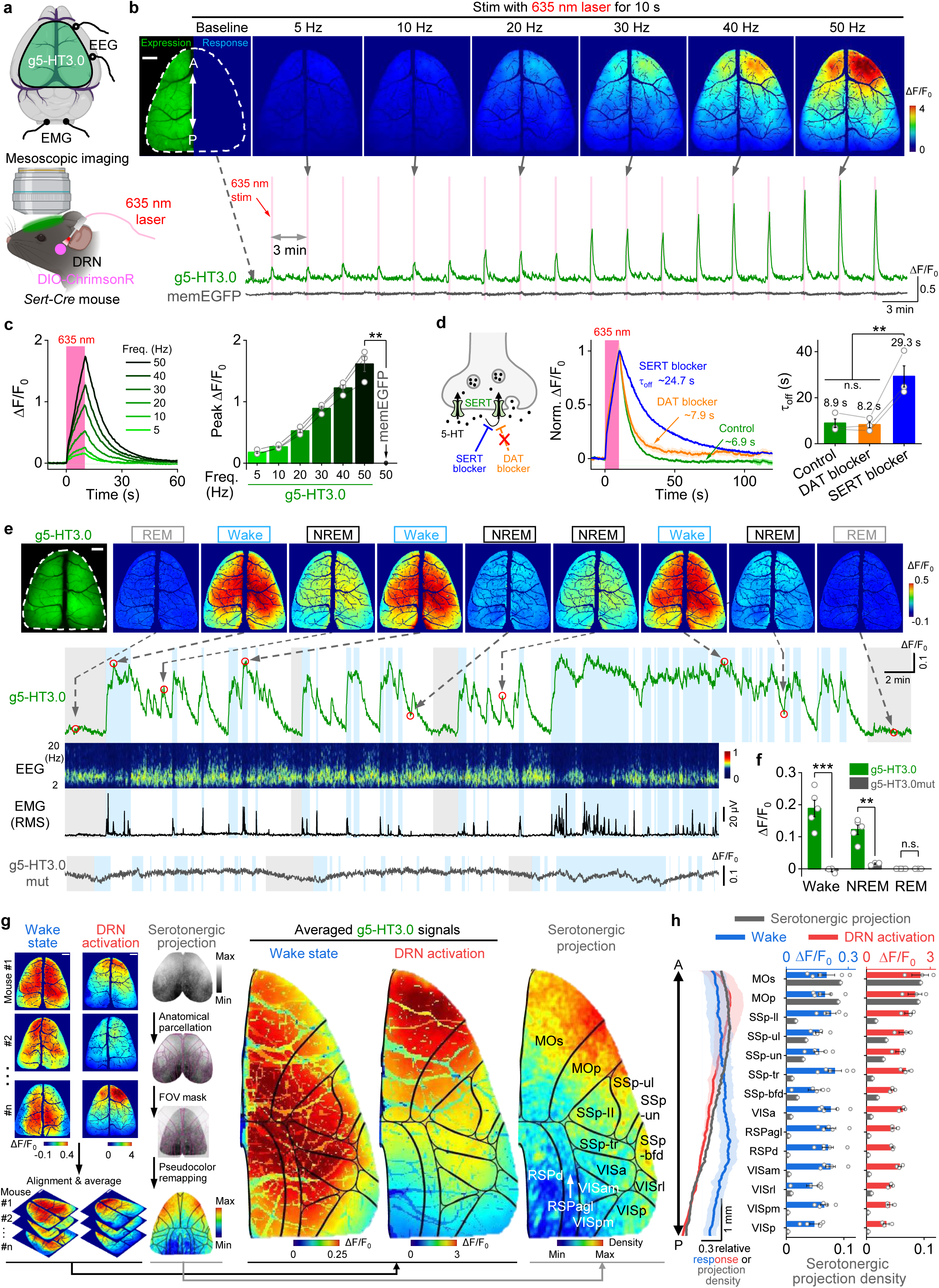
| gGRAB_5-HT3.0_ reveals 5-HT release in mouse dorsal cortex *in vivo* by mesoscopic imaging. **a**, Schematic depicting the setup of mesoscopic imaging. **b**, Representative images showing the cortical g5-HT3.0 expression and response to optical stimulation in the DRN with incremental frequencies (top). Representative traces of g5-HT3.0 and a negative control memEGFP (bottom). The dashed white outline indicates the ROI. **c**, Representative ΔF/F_0_ traces of g5-HT3.0 (left) and group data of peak response (right) with increased frequencies of 635 nm laser. *n* = 3 mice for each group. (Two-tailed Student’s *t*-tests, *P* = 8.48×10^-3^ for g5-HT3.0 versus memEGFP under 50 Hz stimulation.) **d**, Schematic illustrating the effect of SERT blocker and DAT blocker on extracellular 5-HT level (left). Representative ΔF/F_0_ traces of g5-HT3.0 (middle) and summary data of decay kinetics (right) during 50 Hz 10 s stimulation after treatment with indicated compounds. (One-way repeated measures ANOVA followed by Tukey’s multiple-comparison tests, *F* = 28.9, *P* = 4.18×10^-3^, post hoc test: *P* = 0.98 for DAT blocker versus control, 6.45×10^-3^ for SERT blocker versus control and 5.72×10^-3^ for SERT blocker versus DAT blocker.) **e**, Representative fluorescence and pseudocolor images of g5-HT3.0 during the sleep-wake cycle (top). Representative traces of g5-HT3.0 response, EEG, EMG (by root mean square, RMS) and g5-HT3.0mut response in the dorsal cortex during the sleep-wake cycle (bottom). The dashed white outline in the top left image indicates the ROI. Dashed arrows and red circles indicate the timepoint of frames shown at the top. Gray shading, REM sleep; light blue shading, wake state. **f**, Group data of g5-HT3.0 and g5-HT3.0mut responses in mice during the awake state, NREM and REM sleep. *n* = 5 mice for g5-HT3.0 and 3 mice for g5-HT3.0mut. (Two-way repeated measures ANOVA followed by Tukey’s multiple-comparison tests for g5-HT3.0 and g5-HT3.0mut, *P* = 5.77×10^-6^, 1.89×10^-3^ and 1 during the wake, NREM and REM sleep state, respectively.) **g**, Snapshots of g5-HT3.0 responses in different mice in the awake state and DRN activation, and serotonergic projection map modified from Allen Brain (left). Averaged pseudocolor images of g5-HT3.0 responses under indicated conditions (middle) and serotonergic projection map overlaid with black outlines aligned to the Allen Mouse Brain CCF (right). *n* = 5 and 3 mice for the awake state and DRN activation group, respectively. **h**, Averaged relative responses of g5-HT3.0 and serotonergic projection density along the anterior-to-posterior (AP) axis (left) and summary of g5-HT3.0 signals or serotonergic projection density in different cortex regions. All scale bar, 1 mm. Data are shown as mean ± SEM in **c**,**d**,**f**,**h**, with the error bars or shaded regions indicating the SEM, **P < 0.01, ***P < 0.001, n.s., not significant.

Having shown that g5-HT3.0 can reliably detect 5-HT release in the cortex in response to optogenetic stimulation, we then used this sensor to measure physiologically relevant 5-HT dynamics during the sleep-wake cycle using mesoscopic imaging combined with simultaneous EEG and EMG recordings. Similar to our results measured in the subcortical BF (Extended Data Fig. 7), we found that the g5-HT3.0 signal in the dorsal cortex was highest during the wake state, followed by the NREM and REM states, with visible oscillations during NREM sleep. In addition, we found no change in fluorescence through the sleep-wake cycle using the 5-HT insensitive g5-HT3.0mut sensor (Fig. 4e,f).

Taking advantage of our ability to visualize the entire dorsal cortex using mesoscopic imaging, we segmented the dorsal cortex into various brain regions based on the Allen Common Coordinate Framework v3 (CCFv3) atlas^49^ and analyzed the spatial distribution of the 5-HT signals during optogenetic stimulation and during the sleep-wake cycle. We found that the 5-HT signals measured in different brain regions were relatively spatially homogenous and temporally synchronized during the sleep-wake cycle (Fig. 4g and Extended Data Fig. 8h), reminiscent of our previous results recorded in subcortical regions, including the orbital frontal cortex and the bed nucleus of the stria terminalis^17^. In contrast, when we optogenetically stimulated DRN serotonergic neurons, the 5-HT signals had a graded pattern, decreasing along the anterior-to-posterior axis (Fig. 4g,h); interestingly, this pattern was consistent with the anatomically heterogeneous density of serotonergic projections throughout the cortex^44^.

Taken together, these results demonstrate that our next-generation g5-HT3.0 sensor is sufficiently sensitive to monitor 5-HT release *in vivo* with high spatiotemporal resolution, revealing key differences between specific brain regions.

### Dual-color *in vivo* imaging reveals cortex-wide neurochemical waves during seizure activity

The serotonergic system has been suggested to protect the CNS from epileptiform activity^50–54^, which is characterized by excessive and hypersynchronous neuronal firing. However, little is known regarding the spatiotemporal dynamics of 5-HT release during and after seizure activity, let alone the relationship between 5-HT and other seizure-related signals such as calcium^55, 56^ and endocannabinoid (eCB) levels^57^. Therefore, we performed dual-color mesoscopic imaging of g5-HT3.0 together with jRGECO1a (ref.^24^) (to measure both 5-HT and calcium) or r5-HT1.0 together with eCB2.0 (ref.^57^) (to measure both 5-HT and eCBs) in the mouse dorsal cortex, while simultaneously performing EEG recording to identify seizures induced by an injection of the glutamate receptor agonist kainic acid (KA)^58^ (Fig. 5a). Similar to previous reports^57, 59^, we observed an increase in Ca^2+^ during the KA-induced seizure, followed by a spreading wave of Ca^2+^ with a larger magnitude. In the same mouse, we observed a spreading wave of 5-HT, reported by g5-HT3.0 signals, that closely followed the Ca^2+^ wave (Fig. 5b,c,f, Extended Data Fig. 9a,b and Supplementary Video 1). The waves reported by g5-HT3.0 and jRGECO1a originated in approximately the same location and propagated with similar speed (at ∼76 μm/s and ∼83 μm/s, respectively) and in the same direction, primarily from the lateral cortex to the medial region (Fig. 5g,h). As a negative control, seizure activity had no effect on the signal measured using g5-HT3.0mut (Fig. 5c,f and Extended Data Fig. 9a,b).

**Fig. 5.**
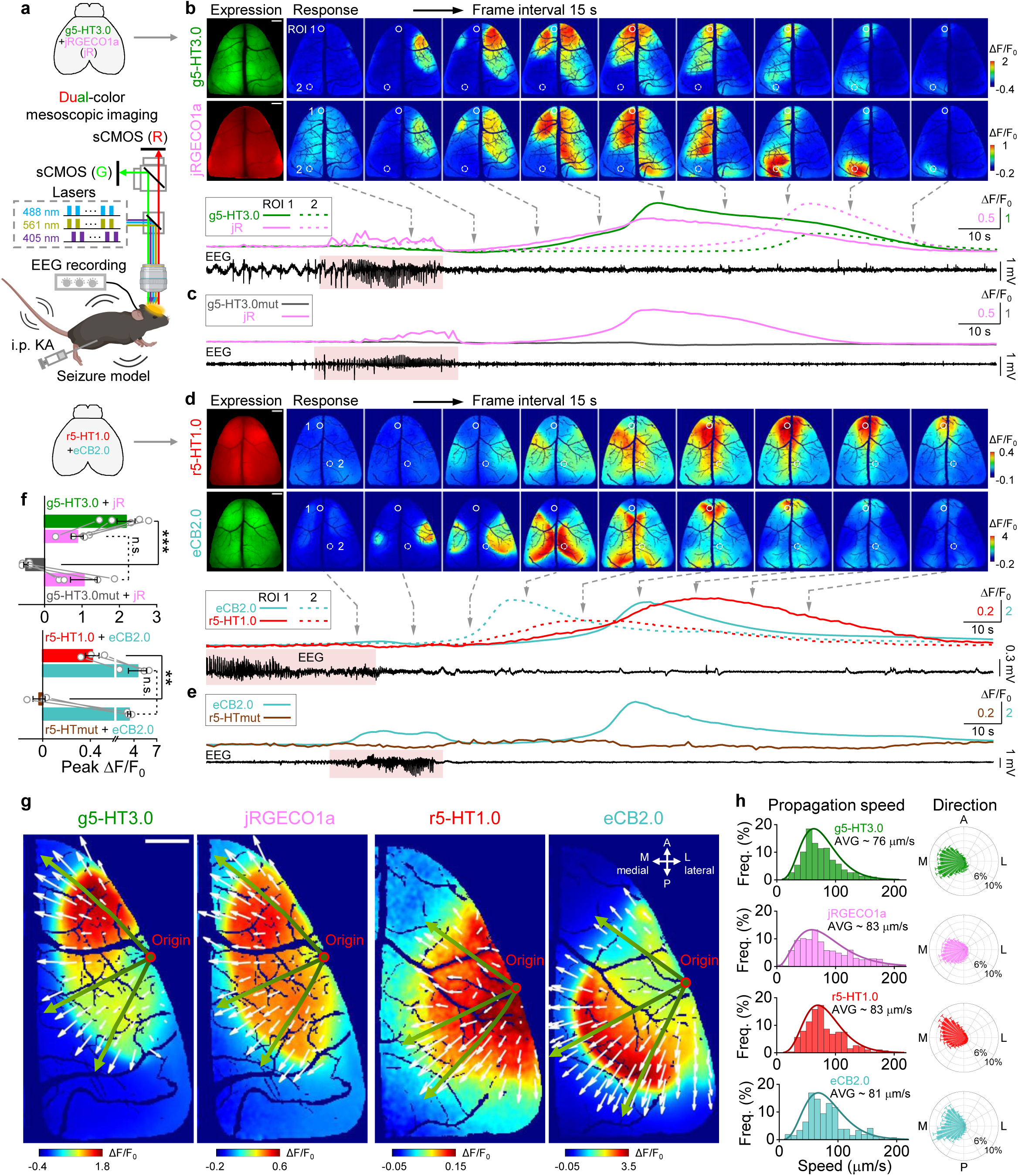
| Dual-color imaging of cortex-wide neurochemical waves during seizures. **a**, Schematic depicting the setup of dual-color mesoscopic imaging in a KA-induced seizure model. **b**, Representative images and ΔF/F_0_ traces of g5-HT3.0 and jRGECO1a during seizures. Two ROIs (500 μm in diameter) are labeled; ROI 1 (the white circle) and ROI 2 (the white dashed circle) show the maximum response regions of g5-HT3.0 and jRGECO1a, respectively. The solid and dashed lines in traces correspond to ROI 1 and ROI 2, respectively. The red shading in the EEG trace indicates the epileptic discharges. **c**, Representative ΔF/F_0_ traces of g5-HT3.0mut and jRGECO1a during seizures, similar to **b**, and images are showed in Extended Data Fig. 9b. **d**, Representative images and ΔF/F_0_ traces of r5-HT1.0 and eCB2.0 during seizures, similar to **b**, except that ROI 1 (the white circle) and ROI 2 (the white dashed circle) show the maximum response regions of r5-HT1.0 and eCB2.0, respectively. **e**, Representative traces of r5-HTmut and eCB2.0 signals during seizures, similar to **d**, and images are showed in Extended Data Fig. 9d. **f**, Group summary of different sensors’ peak responses. *n* = 5 mice for the group co-expressing g5-HT3.0 and jRGECO1a, *n* = 4 for g5-HT3.0mut and jRGECO1a, *n* = 3 for r5-HT1.0 and eCB2.0, *n* = 3 for r5-HTmut and eCB2.0. (Two-tailed Student’s *t*-tests, *P* = 2.36×10^-4^ for g5-HT3.0 versus g5-HT3.0mut, *P* = 0.64 for jRGECO1a between two groups; *P* = 4.41×10^-3^ for r5-HT1.0 versus r5-HTmut, *P* = 0.45 for eCB2.0 between two groups.) **g**, Representative images showing the wave propagation detected by indicated sensors. The red circle indicates the origin of waves; small white arrows indicate the wave-propagating velocity vector; green lines with arrow indicate example propagating trajectories. L, lateral, M, medial, A, anterior, P, posterior. **h**, Probability distributions of wave-propagating speed and direction calculated by indicated sensors. Scale bar in all images, 1 mm. Data are shown as mean ± SEM in **f**,**h**, with the error bars indicating the SEM, \*\**P* < 0.01, \*\*\**P* < 0.001, n.s., not significant.

Finally, we obtained similar results in mice co-expressing r5-HT1.0 and eCB2.0, with a propagating wave of 5-HT release following the eCB wave. Moreover, the waves reported by r5-HT1.0 and eCB2.0 originated in approximately the same location, propagated at similar speed (∼83 μm/s and ∼81 μm/s, respectively) and in the same direction (Fig. 5d,f– h and Supplementary Video 2). As above, seizure activity had no effect on the signal measured using r5-HTmut (Fig. 5e,f and Extended Data Fig. 9c,d).

Taken together, these results demonstrate that our g5-HT3.0 and r5-HT1.0 sensors can reliably report 5-HT release *in vivo* with high sensitivity, specificity, and spatiotemporal resolution both under physiological conditions and during seizure activity.

## Discussion

Here, we report the development, optimization, characterization, and *in vivo* application of a series of genetically encoded red and green fluorescent 5-HT sensors. These 5-HT sensors have high specificity, sensitivity, and spatiotemporal resolution, as well as rapid kinetics. More importantly, they do not couple to downstream signal pathways. Both the g5-HT3.0 and r5-HT1.0 sensors reliably reported time-locked 5-HT release induced by optogenetic activation of DRN serotonergic neurons, as well as changes in 5-HT levels during the sleep-wake cycle. Moreover, using dual-color mesoscopic imaging, we found that seizure activity triggers a cortex-wide 5-HT wave that follows a wave of Ca^2+^ and eCBs. GRAB sensors consist of a ligand-binding module and a fluorescent reporting module^60^.

Our ability to efficiently develop new GRAB_5-HT_ sensors was facilitated by our transplantation strategy and our selection of the 5-HTR4 receptor as the scaffold. First, we used the transplantation strategy to capitalize on previously optimized reporting modules and linkers in existing sensors. Next, we chose the best-performing scaffold (5-HTR4) from among the various 5-HT receptor subtypes as the ligand-binding module, thereby accelerating the optimization processes. A similar strategy can be used in order to accelerate the development of other GPCR-based sensors.

Our next-generation green fluorescent GRAB_5-HT_ sensors have three advantages over previously reported genetically encoded fluorescent 5-HT sensors. First, g5-HT3.0 is significantly more sensitive than other GPCR- and PBP-based sensors due to its larger response and brightness, making it more suitable for both *in vitro* and *in vivo* applications. Second, our high-affinity g5-HT3.0 (EC_50_: ∼150 nM) and medium-affinity g5-HT2m (EC_50_: ∼1.1 µM) sensors fill the critical gap in measuring intermediate concentrations of 5-HT. Thus, together with previous sensors such as g5-HT1.0 (EC_50_: ∼20 nM) and iSeroSnFR (EC_50_: >300 μM), we now possess a powerful toolbox covering a wide range of physiological and pathological 5-HT concentrations^61–63^. Third, these sensors have rapid kinetics and are suitable for tracking transient 5-HT release in real time.

More importantly, our all-new red fluorescent 5-HT sensor expands the spectral profile of genetically encoded 5-HT sensors and is suitable for dual-color imaging when combined with green fluorescent sensors. Here, we simultaneously recorded red and green fluorescence using r5-HT1.0 and axon-targeted GCaMP6s, respectively, in the mouse basal forebrain during the sleep-wake cycle and found that the two signals were closely correlated, consistent with Ca^2+^-dependent rapid 5-HT release. In the future, further expanding the spectra of 5-HT sensors (for example, to include the far-red and near-infrared spectra) will provide the ability to simultaneously monitor multiple signals in addition to 5-HT.

Using mesoscopic imaging, we found that 5-HT release in the mouse dorsal cortex differs under different behavioral conditions. A previous study using a calcium sensor suggested that 5-HT release might differ between the orbitofrontal cortex and central amygdala in response to rewarding and aversive stimuli, but the 5-HT release was not measured directly^37^. Recently, using g5-HT1.0, we showed that 5-HT release is highly synchronized between different brain regions during NREM sleep^17^. Here, we expanded on this finding using mesoscopic imaging of g5-HT3.0 to image more brain regions in the mouse dorsal cortex and found relatively homogenous release throughout the dorsal cortex during the sleep-wake cycle. Moreover, we found a graded pattern of 5-HT release in the mouse dorsal cortex evoked by optogenetic activation of the DRN, similar to the pattern of serotonergic projections. These results suggest that although the potential 5-HT release is spatially heterogenous throughout the dorsal cortex, this release may be regulated on a global scale during the sleep-wake cycle, leading to relatively homogenous 5-HT release under these conditions. Furthermore, using a KA-induced seizure model, we found that 5-HT release propagates as a wave across the cortex, a result made possible due to the advantages of our new 5-HT sensors compared to traditional methods used to measure 5-HT. Indeed, a previous study using microdialysis found increased 5-HT levels after seizures in rats, but the results had relatively poor temporal resolution and lacked spatial resolution^54^. Interestingly, we further found that the seizure-induced 5-HT waves were spatially correlated—but lagged behind—Ca^2+^ and eCB waves, consistent with the idea that serotonergic activity may protect against neuronal hyperactivity^51, 53, 64^.

In summary, our new 5-HT sensors can be used to monitor 5-HT release both *in vitro* and *in vivo*, with high sensitivity and spatiotemporal resolution. Thus, when combined with advanced imaging techniques, these new 5-HT sensors provide a robust toolbox to study the serotonergic system in both health and disease.

## Methods

### Molecular biology

DNA fragments were amplified by PCR using primers (RuiBiotech) with 25–30-bp overlap and used to generate plasmids via the Gibson assembly method^65^ using T5 exonuclease (New England Biolabs), Phusion DNA polymerase (Thermo Fisher Scientific), and Taq ligase (iCloning). Plasmid sequences were verified by Sanger sequencing (RuiBiotech). For the replacement site screening, cDNAs encoding 12 different 5-HTR subtypes (5-HTR_1A_, 5-HTR_1B_, 5-HTR_1D_, 5-HTR_1E_, 5-HTR_1F_, 5-HTR_2A_, 5-HTR_2B_, 5-HTR_2C_, 5-HTR_4_, 5-HTR_5A_, 5-HTR_6_, and 5-HTR_7_) were cloned by PCR amplification of the full-length human GPCR cDNA library (hORFeome database 8.1) or the PRESTO-Tango GPCR Ki^t36^ (Addgene Kit #1000000068). To optimize the 5-HT sensors, cDNAs encoding candidate sensors were cloned into the pDisplay vector (Invitrogen) with an IgK leader sequence in the sensor upstream, and either IRES-mCherry-CAAX (for green fluorescent 5-HT sensors) or IRES-EGFP-CAAX (for red fluorescent 5-HT sensors) was fused downstream of the sensor to calibrate the membrane signal. Site-directed mutagenesis was performed using primers containing randomized NNB codons (48 codons in total, encoding 20 possible amino acids) or defined codons at the target sites. For expression in cultured neurons, the sensors were cloned into the pAAV vector under the control of the hSyn promoter. To generate stable cell lines for measuring the excitation/emission spectra, sequences encoding various 5-HT sensors were cloned into a vector called pPacific, containing a 3’ terminal repeat, IRES, the puromycin gene, and a 5’ terminal repeat. Two mutations (S103P and S509G) were introduced into pCS7-PiggyBAC to generate hyperactive piggyBac transposase (ViewSolid Biotech)^66^. To measure downstream coupling using the Tango assay, DNA encoding various GRAB_5-HT_ sensors or wild-type 5-HTR4 was cloned into the pTango vector^36^. For the luciferase complementation assay, the β_2_AR gene in the β2AR-Smbit construct^34^ was replaced with the indicated GRAB_5-HT_ sensors or wild-type 5-HTR4; LgBit-mGs was a generous gift from Nevin A. Lambert (Augusta University).

### Cell lines

HEK293T cells were purchased from ATCC (CRL-3216) and verified based on their morphology and growth rate. Stable cell lines expressing different GRAB_5-HT_ sensors were generated by co-transfecting HEK293T cells with the pPacific plasmids encoding sensors and the pCS7-PiggyBAC plasmid encoding the transposase^66^. Cells expressing the desired genes were selected using 2 μg/ml puromycin (Sigma). An HTLA cell line stably expressing a tTA-dependent luciferase reporter and the β-arrestin2-TEV fusion gene used in the Tango assay^36^ was a generous gift from Bryan L. Roth (University of North Carolina Chapel Hill). All cell lines were cultured at 37°C in 5% CO_2_ in DMEM (Biological Industries) supplemented with 10% (v/v) fetal bovine serum (GIBCO) and 1% penicillin-streptomycin (GIBCO).

### Primary cultures

Rat cortical neurons were prepared using postnatal day 0 (P0) Sprague-Dawley rat pups (both sexes) purchased from Beijing Vital River. The cerebral cortex was dissected, and neurons were dissociated using 0.25% trypsin-EDTA (GIBCO), plated onto 12-mm glass coverslips coated with poly-D-lysine (Sigma-Aldrich), and cultured in neurobasal medium (GIBCO) containing 2% B-27 supplement (GIBCO), 1% GlutaMax (GIBCO), and 1% penicillin-streptomycin (GIBCO) at 37°C in humidified air containing 5% CO_2_.

### Animals

All procedures involving animals were performed using protocols approved by the Animal Care and Use Committee at Peking University. *Sert-Cre* mice were generously provided by Yi Rao at Peking University. All mice were group-housed or pair-housed in a temperature-controlled room (18–23°C) with a 12-h/12-h light/dark cycle, with food and water available ad libitum.

### Cell transfection and imaging

HEK293T cells were plated either on 12-mm glass coverslips in 24-well plates or 96-well plates without coverslips and grown to ∼70% confluence for transfection with PEI (1 μg plasmid and 3 μg PEI per well in 24-well plates or 300 ng plasmids and 900 ng PEI per well in 96-well plates); the medium was replaced after 4–6 h, and the cells were used for imaging 24–48 h after transfection. After 5–9 d of *in vitro* culture, rat cortical neurons were infected with AAVs expressing the following 5-HT sensors: g5-HT3.0 (5.72×10^12^ vg/ml, BrainVTA), g5-HT2m (2.53×10^12^ vg/ml, BrainVTA), g5-HT2h (2.67×10^12^ vg/ml, BrainVTA), g5-HT1.0 (AAV9-hSyn-tTA, 2.39×10^13^ vg/ml, and AAV9-TRE-5-HT1.0, 3.81×10^13^ vg/ml, Vigene Biosciences), iSeroSnFR (2.76×10^12^ vg/ml, BrainVTA), PsychLight2 (3.07×10^12^ vg/ml, BrainVTA), g5-HT3.0mut (6.16×10^13^ vg/ml, Vigene Biosciences), r5-HT1.0 (9.44×10^13^ vg/ml, Vigene Biosciences), or r5-HTmut (1.17×10^14^ vg/ml, Vigene Biosciences). To compare the performance of various sensors, the volume of each AAV was adjusted by titer to apply the same number of AAV particles.

Before imaging, the culture medium was replaced with Tyrode’s solution consisting of (in mM): 150 NaCl, 4 KCl, 2 MgCl_2_, 2 CaCl_2_, 10 HEPES, and 10 glucose (pH 7.4).

Cells were imaged using an inverted Ti-E A1 confocal microscope (Nikon) or an Opera Phenix high-content screening system (PerkinElmer). The confocal microscope was equipped with a 10x/0.45 NA (numerical aperture) objective, a 20x/0.75 NA objective, a 40x/1.35 NA oil-immersion objective, a 488-nm laser, and a 561-nm laser; the GFP signal was collected using a 525/50-nm emission filter combined with the 488-nm laser, and the RFP signal was collected using a 595/50-nm emission filter combined with the 561-nm laser. The Opera Phenix system was equipped with a 20x/0.4 NA objective, a 40x/1.1 NA water-immersion objective, a 488-nm laser, and a 561-nm laser; the GFP and RFP signals were collected using a 525/50-nm and 600/30-nm emission filter, respectively. The fluorescence signals produced by the green and red fluorescent GRAB_5-HT_ sensors were calibrated using mCherry (the GFP/RFP ratio) or EGFP (the RFP/GFP ratio), respectively.

To measure the sensor responses induced by various chemicals, solutions containing the indicated concentrations of 5-HT and/or other chemicals were administered to the cells via a custom-made perfusion system or via bath application. To measure the sensors’ kinetics, a glass pipette was positioned in close proximity to cells expressing the sensors, and the fluorescence signal was measured using confocal high-speed line scanning mode with a scanning speed of 1024 Hz. To measure the on-rate constant (τ_on_), 100 μM 5-HT was puffed from the pipette, and the increased trace in fluorescence was fitted with a single-exponential function; to measure the off-rate constant (τ_off_), 100 μM RS 23597-190 was puffed on cells bathed in 10 μM 5-HT, and the decreased trace in fluorescence was fitted with a single-exponential function. To test for blue light-mediated photoactivation of the red fluorescent sensors, a 488-nm laser lasting for 1 s was utilized (power: ∼210 μW, intensity: ∼0.4 W/cm^2^).

### Spectra measurements

For 1-photon spectra, HEK293T cells stably expressing g5-HT3.0, g5-HT2m, g5-HT2h, or r5-HT1.0 were harvested and transferred to a 384-well plate in the absence or presence of 10 μM 5-HT. Excitation and emission spectra were measured at 5-nm increments with a 20-nm bandwidth using a Safire2 multi-mode plate reader (Tecan). To obtain background for subtraction, control cells (not expressing a sensor) were prepared to the same density as the cells that expressed sensors, and were measured using the same protocol.

For 2-photon spectra, HEK293T cells expressing g5-HT3.0, g5-HT2m, g5-HT2h, or r5-HT1.0) were cultured on 12-mm coverslips and placed in an imaging chamber under a 2-photon microscope. For green fluorescent sensors, 2-photon excitation spectra were measured at 10-nm increments ranging from 690 to 1030 nm using a 2-photon microscope (Bruker) equipped with a 20x/1.0 NA water-immersion objective (Olympus) and an InSight X3 tunable laser (Spectra-Physics). For the red fluorescent sensor, 2-photon excitation spectra were measured at 10-nm increments ranging from 820 to 1300 nm using an A1R MP+ multiphoton microscope (Nikon) equipped with a 25x/1.1 NA objective (Nikon) and a Chameleon Discovery tunable laser (Coherent). Laser power was calibrated for various wavelengths.

### Luciferase complementation assay

The luciferase complementation assay was performed as previously described^34^. In brief, 24–48 h after transfection, the cells were washed with PBS, dissociated using a cell scraper, resuspended in PBS, transferred to opaque 96-well plates containing 5 μM furimazine (NanoLuc Luciferase Assay, Promega), and bathed in 5-HT at various concentrations (ranging from 0.1 nM to 1 mM). After incubation for 10 minutes in the dark, luminescence was measured using a VICTOR X5 multilabel plate reader (PerkinElmer).

### Tango assay

A reporter cell line called HTLA, stably expressing a tTA-dependent luciferase reporter and a β-arrestin2-TEV fusion gene, was transfected with pTango vectors to express GRAB_5-HT_ sensors or wild-type 5-HTR4. After culturing for 24 h in 6-well plates, the cells were transferred to 96-well plates and bathed with 5-HT at varying concentrations (ranging from 0.01 nM to 100 μM). The cells were then cultured for 12 h to allow the expression of tTA-dependent luciferase. Bright-Glo reagent (Fluc Luciferase Assay System, Promega) was added to a final concentration of 5 μM, and luminescence was measured using a VICTOR X5 multilabel plate reader (PerkinElmer).

### Fiber photometry recording of 5-HT release *in vivo*

To express the red fluorescent 5-HT sensors in the BF, adult *Sert-Cre* mice were anesthetized with 1.5% isoflurane and placed on a stereotaxic frame (RWD Life Science). AAV expressing hSyn-r5-HT1.0 (9.44×10^13^ vg/ml, Vigene Biosciences) or hSyn-r5-HTmut (1.14×10^14^ vg/ml, Vigene Biosciences) was injected (400 nl per site) via a glass pipette using a micro-syringe pump (Nanoliter 2000 Injector, World Precision Instruments) at the following coordinates: AP: 0 mm relative to Bregma; ML: +1.5 mm; DV: 4.6 mm below the dura. For optical activation of the DRN, 400 nl AAV9-EF1a-DIO-hChR2(H134R)-EYFP (5.54×10^13^ vg/ml, BrainVTA) was injected into the DRN at the following coordinates: AP: -

4.1 mm relative to Bregma; ML: +1.1 mm; depth: 2.9 mm below the dura; at a 20° ML angle). Two optical fiber cannulas (200 μm, 0.37 NA, Inper, Zhejiang, China) were then implanted; one cannula was implanted 0.1 mm above the virus injection site in the BF to record the 5-HT sensors, and the other cannula was implanted 0.3 mm above the virus injection site in the DRN for optically activating ChR2. The optical fibers were affixed to the skull surface using dental cement. The fiber photometry system (Inper Tech, Zhejiang, China) was used to record the fluorescence signals in freely moving mice. Yellow light-emitting diode (LED) light was bandpass filtered (561/10 nm), reflected by a dichroic mirror (495 nm), and then focused using a 20x objective lens (Olympus). An optical fiber was used to guide the light between the commutator and the implanted optical fiber cannulas. The excitation light emitted by the LED was adjusted to 20–30 μW and delivered at 10 Hz with a 20-ms pulse duration. The optical signals were then collected through the optical fibers. Red fluorescence was bandpass filtered (520/20 nm and 595/30 nm) and collected using an sCMOS camera. To induce ChR2-mediated 5-HT release, pulse trains (10-ms pulses at 50 Hz for 1 s, 5 s, or 10 s) were delivered to the DRN using a 488-nm laser at 20 mW with a 5-min inter-stimulus interval. To test the effects of fluoxetine on the ChR2-induced responses, three optical stimulation trains were applied at a 5-min interval. Then, 10 mg/kg fluoxetine was administered via i.p. injection; 30 min after injection, three optical stimulation trains were applied.

The current output from the photomultiplier tube was converted to a voltage signal using a model 1700 differential amplifier (A-M Systems) and passed through a low-pass filter. The analog voltage signals were then digitized using an acquisition card (National Instruments). To minimize autofluorescence of the optical fibers, the recording fibers were photobleached using a high-power LED before recording. Background autofluorescence was recorded and subtracted from the recorded signals in the subsequent analysis.

The photometry data were analyzed using a custom program written in MATLAB. To calculate ΔF/F_0_ during the optogenetics experiments, a baseline was measured before optical stimulation.

### Fiber photometry recording of 5-HT dynamics during the sleep-wake cycle

Adult wild-type C57BL/6 mice and *Sert-Cre* mice were anesthetized with isoflurane and placed on a stereotaxic frame (RWD Life Science). In Extended Data Fig. 7a–c, AAV expressing CAG-g5-HT1.0 (3.16×10^12^ vg/ml) was injected into the BF in one hemisphere and AAV expressing CAG-g5-HT3.0 (3.16×10^12^ vg/ml) was injected into the BF in another hemisphere (400 nl per site) using the coordinates described above. In Extended Data Fig. 7d–f, AAV expressing hSyn-PsychLight2 (3.07×10^12^ vg/ml) and hSyn-g5-HT3.0 (5.72×10^12^ vg/ml, diluted to 3.07×10^12^ vg/ml) were injected into the bilateral BF (400 nl per site), respectively. In Extended Data Fig. 7g–i, AAV expressing CAG-iSeroSnFR (2.15×10^12^ vg/ml) and CAG-5-HT3.0 (3.16×10^12^ vg/ml, diluted to 2.15×10^12^ vg/ml) were injected into the bilateral BF (400 nl per site), respectively. In Extended Data Fig. 7, all AAVs were produced by BrainVTA and mice were wild-type. In Fig. 3f–g, AAV expressing hSyn-r5-HT1.0 (9.44×10^13^ vg/ml, Vigene Biosciences) or hSyn-r5-HTmut (1.14×10^14^ vg/ml, Vigene Biosciences) was injected into the BF, and EF1α-DIO-axon-GCaMP6s (5.71×10^12^ vg/ml, BrainVTA) was injected into the DRN (400 nl per site) of *Sert-Cre* mice. An optical fiber cannula (200 μm, 0.37 NA, Inper, Zhejiang, China) was implanted 0.1 mm above the virus injection sites in BF for recording the signals of 5-HT sensors and calcium sensor.

To record the animal’s sleep-wake state, we attached and fixed custom-made EEG and EMG electrodes to the skull via a microconnector. EEG electrodes were implanted into craniotomy holes situated over the frontal cortex and visual cortex, and EMG wires were placed bilaterally in the neck musculature. The microconnector was attached to the skull using glue and a thick layer of dental cement. After surgery, the mice were allowed to recover for at least 2 weeks.

The same fiber photometry system (Inper Tech, Zhejiang, China) was used to record the fluorescence signals in freely moving mice during the sleep-wake cycle. In Extended Data Fig. 7, a 10-Hz 470/10-nm filtered light (20-30 µW) was used to excite green fluorescent 5-HT sensors, and a 520/20-nm and 595/30 nm dual-band bandpass filter was used to collect the fluorescence signals. In Fig. 3f,g, a 10-Hz 470/10-nm filtered light (20–30 µW) was used to excite the green fluorescent calcium sensor, and a 561/10-nm filtered light (20–30 µW) was used to excite the red fluorescent 5-HT sensors. Fluorescence signals were collected using a dual-band bandpass filter (520/20 nm and 595/30 nm), with excitation light delivered as 20-ms pulses at 10 Hz.

Photometry data were analyzed using a custom MATLAB program. To calculate the *z*-score during the sleep-wake cycle, baseline values were measured during a period of REM sleep in which no apparent fluctuations were observed.

### Polysomnographic recording and analysis

The animal’s sleep-wake state was determined using the EEG and EMG recordings. The EEG and EMG signals were amplified (NL104A, Digitimer), filtered (NL125/6, Digitimer) at 0.5–100 Hz (EEG) and 30–500 Hz (EMG), and then digitized using a Power1401 digitizer (Cambridge Electronic Design Ltd.). The Spike2 software program (Cambridge Electronic Design Ltd.) was used for recording with a sampling rate of 1000 Hz. The animal’s sleep-wake state was classified semi-automatically in 4-s epochs using AccuSleep^67^ and then validated manually using a custom-made MATLAB GUI. The wake state was defined as desynchronized EEG activity combined with high EMG activity. NREM sleep was defined as synchronized EEG activity with high-amplitude delta activity (0.5–4 Hz) and low EMG activity. REM sleep was defined as high-power theta frequencies (6–9 Hz) combined with low EMG activity.

### Mesoscopic *in vivo* imaging

To express the sensors throughout the cortex, we injected the indicated AAVs into the transverse sinus as described previously^45^. In detail, P0–P1 C57BL/6 mouse pups were removed from their home cages, placed on a warm pad, anesthetized on ice for 2–3 min, and fixed on an ice cooled metal plate. Two small incisions were then made over the transverse sinuses for AAV injection using a glass pipette. The pups were injected bilaterally with the following pairs of AAVs (6 μl total volume): AAV9-hSyn-5-HT3.0 (8.67×10^13^ vg/ml, Vigene Biosciences) and AAV9-hSyn-GAP43-jRGECO1a (1.47×10^13^ vg/ml, OBiO); AAV9-hSyn-r5-HT1.0 (9.44×10^13^ vg/ml, Vigene Biosciences) and AAV9-hSyn-eCB2.0 (9.11×10^13^ vg/ml, Vigene Biosciences); AAV9-hSyn-5-HT3.0mut (6.16×10^13^ vg/ml, Vigene Biosciences) and AAV9-hSyn-GAP43-jRGECO1a; or AAV9-hSyn-r5-HTmut (1.17×10^14^ vg/ml, Vigene Biosciences) and AAV9-hSyn-eCB2.0. The AAVs were injected at a rate of 1.2 μl/min, and the pipette was left in the sinus for at least 30 s. After injection, the incisions were sealed with Vetbond glue (3M Animal Care Products) and the pups were placed on a warm pad for recovery. After recovery, the pups were gently rubbed with bedding and returned to their home cage.

About eight weeks after AAV injection, surgery was performed for implanting the imaging window and the EEG and EMG electrodes. Anesthesia was induced with an i.p. injection of 2,2,2-tribromoethanol (Avertin, 500 mg/kg, Sigma-Aldrich) and maintained using inhalation with 1% isoflurane. The mouse was fixed in a stereotaxic frame, 2% lidocaine hydrochloride was injected under the scalp, and the eyes were covered with erythromycin ophthalmic ointment for protection. Part of the scalp above the skull and the underlying muscles were removed and cleaned carefully to expose the skull. Most of the skull above the dorsal cortex was then carefully removed and replaced with a flat custom-made coverslip (D-shape, ∼8 mm × 8 mm) to create an optical window. EEG and EMG electrodes were implanted and fixed as described above. After surgery, the mice were returned to their home cage for at least 7 d to recover, and then fixed to the base for over 3 d to habituate before imaging until the mouse can fall asleep (especially REM sleep) within the first 3 h. To optically activate the DRN, we used *Sert-Cre* mice as described above, except before the surgery, 300 nl AAV9-EF1a-DIO-ChrimsonR-iP2A-Halotag9-V5 (6.81×10^13^ vg/ml, Vigene Biosciences) was injected into the DRN using the following coordinates: AP:

−6.1 mm relative to Bregma; ML: 0 mm relative to Bregma; depth: 3 mm below the dura; at a 32° AP angle (to avoid the imaging window, the fiber is inserted forward and down from the back of the interparietal bone). An optical fiber cannula (200 μm, 0.37 NA, Inper, Zhejiang, China) was then implanted 0.2 mm above the virus injection site and affixed to the skull surface using dental cement.

Mesoscopic imaging was performed using a custom-made dual-color macroscope equipped with a 2x/0.5 NA objective lens (Olympus, MVPLAPO2XC), two 1x/0.25 NA tube lenses (Olympus, MVPLAPO1X), and two sCMOS cameras (Andor, Zyla 4.2 Plus, 2,048×2,048 pixels, 16-bit)^68, 69^. Three excitation wavelengths (405 nm, 488 nm, and 561 nm) were generated using a multi-line fiber coupled laser system (Changchun New Industries Optoelectronics Tech. Co., Ltd., RGB-405/488/561/642nm-220mW-CC32594). The emission light was passed through a 567-nm cut-on longpass dichroic mirror (Thorlabs, DMLP567L), then through either a 525/36-nm or 609/34-nm emission filter (Chroma), and finally into the sCMOS cameras (one for the green channel and one for the red channel). Both the excitation laser and the camera imaging were triggered by an Arduino board (Uno) with custom-written programs. For green sensor imaging, a 488-nm laser was used and interleaved with a 405-nm laser; while for dual-color imaging, 488-nm and 561-nm lasers were simultaneously generated and interleaved with a 405-nm laser. Images were acquired using Micro-Manager 2.0 at a resolution of 512 × 512 pixels after 4× pixel binning, and each channel was acquired at either 1 Hz or 5 Hz with 40-ms exposure.

During imaging, the mice were head-fixed to the base and could freely run on a treadmill^70^. An infrared camera equipped with LEDs was placed around the mouse’s head to capture its face and eyes for recording behavioral data. To induce ChrimsonR-mediated 5-HT release in the optogenetics experiment, pulse trains of light (10-ms pulses for 10 s at 5, 10, 20, 30, 40, or 50 Hz) were applied to the DRN using a 635-nm laser at 10 mW with a 3-minute interval, and three trials were performed at each frequency. To test the effects of fluoxetine and GBR 12909 on ChrimsonR-evoked 5-HT release, three trains of optical stimulation (10-ms pulses for 1 s at 20 Hz, 10-ms pulses for 10 s at 20 Hz, or 10-ms pulses for 10 s at 50 Hz) were applied with a 5-min interval, and three trials were conducted for each parameter. Then, 10 mg/kg GBR 12909 was injected i.p., and the same optical stimulation procedure was performed 30 min later. Next, 20 mg/kg fluoxetine was injected i.p., followed by the same optical stimulation procedure 30 min later (see Extended Data Fig. 8a). For the seizure experiments, another infrared camera was hung above the back of the mouse to record its behavioral data. After recording ∼1 h of baseline data, 10 mg/kg KA was injected i.p. to induce seizures. All recordings, including mesoscopic imaging, EEG and EMG recording, optical stimulation trains, and the infrared cameras, were synchronized using a Power1401 acquisition board (Cambridge Electronic Design Ltd.).

### Analysis of mesoscopic imaging data

#### Preprocessing

Raw images acquired by each camera were calibrated for the uniformity of the imaging system, and movement-related artifacts were corrected using the motion-correction algorithm NoRMCorre (ref.^71^). The corrected image stack with a size of 512 × 512 pixels was downsampled by a factor of 0.7 to 359 × 359 pixels for subsequent analysis. For dual-color imaging, the averaged red-channel image was registered to the averaged green-channel image via automatic transformation using the MATLAB function “imregtform” with the “similarity” mode. The same geometric transformation was applied to all red-channel images to register to corresponding green-channel images. The image stack was saved as a binary file to accelerate the input and output of large files (typically >8 GB). To remove pixels belonging to the background and blood vessels (particularly large veins), we generated a mask for further analysis of the pixels. Specifically, the outline of the entire dorsal cortex in the field of view was generated manually, and pixels outside the outline were set as background and excluded from further analysis. The blood vessels were then removed from the image using the machine learning-based ImageJ plugin Trainable Weka Segmentation^72^ (v3.3.2) in order to minimize artifacts caused by the constriction and dilation of blood vessels. The final mask without the background and blood vessel pixels was applied to the image stack for further analysis.

#### Spectral unmixing

For simultaneous dual-color imaging, the bleed-through of fluorescence intensity for each pixel between the green and red channels was removed using linear unmixing based on the spectra of the various sensors and the setup of the microscope system.

#### Hemodynamic correction and response calculation

Hemodynamic changes can affect the absorption of light, resulting in changes in fluorescence^73, 74^. According to the spectra of the sensors, when excited with 405-nm light, the g5-HT3.0, eCB2.0, r5-HT1.0, and jRGECO1a sensors are ligand-insensitive, which can reflect hemodynamic absorption. To correct for hemodynamic artifacts, we performed a pixel-by-pixel correction based on linear regression^75^ of the ligand-dependent signals (excited by 488 nm or 561 nm) against the ligand-independent signals (excited by 405 nm). The baseline images were spatially smoothed using a Gaussian filter (σ=2) and used for the linear regression. Then, for each pixel, the baseline fluorescence intensity of the 405-nm excited channel was regressed onto the 488-nm or 561-nm signal, obtaining regression coefficients for rescaling the 405-nm channel. The rescaled 405-nm signal was subtracted from the 488-nm or 561-nm signal to generate a corrected signal for each pixel. To avoid the corrected signal becoming close to zero or even less than zero, the corrected signal was added to the average rescaled 405-nm channel signal as the final corrected signal for the response calculation. The response of each pixel was calculated using the equation ΔF/F_0_ = (F−F_0_)/F_0_, where F_0_ is defined as the average baseline fluorescence intensity. When analyzing the data obtained during the sleep-wake cycle, the baseline was defined as the REM sleep state, in which the signal had low fluctuations.

#### Parcellation of cortical areas

Based on previous studies^76, 77^, we rigidly registered the average fluorescence image to a 2D projection of the Allen Common Coordinate Framework v3 (CCFv3) using four manually labeled anatomical landmarks, namely the left, center, and right points in the boundary between the anterior cortex and the olfactory bulbs, and the medial point at the base of the retrosplenial cortex. To analyze the time series response in an individual brain region, we averaged the ΔF/F_0_ value for all available pixels within that brain region. To obtain the average response map from multiple mice (see Fig. 4g), we developed a custom code that first register the response image for each individual mouse to the Allen CCFv3, and then averaged the images, keeping only the intersection among all mice. For the analysis of serotonergic projection in the mouse dorsal cortex, the serotonergic projection map was modified from Allen Mouse Brain Connectivity Atlas, connectivity.brain-map.org/projection/experiment/cortical_map/480074702.

### Analysis of propagating waves

#### Peak response calculation

The time series of the images obtained before (∼30 s before the wave originated), during, and after (∼30 s after the wave disappeared) wave propagation was extracted as an event for further analysis. Images taken during the first 20 s (20 frames) were set as the event baseline. The event response image was spatially filtered, and each pixel during the event was then corrected to set the average response of the event baseline to zero. The peak response site was automatically found by a circle with a 500-μm diameter across the event, and its average value was defined as the peak response of the event.

#### Identification of wave directions using optical flow analysis

To determine the direction of the waves, we adopted an optical flow method for automatically detecting the wave directions based on the NeuroPatt toolbox^78^. In detail, the corrected response image stack was smoothed over time, downsampled in size by a factor of 0.2, and normalized to the maximum response for each pixel. The phase velocity fields were then calculated using the “opticalFlowHS” MATLAB function (smoothness parameter α = 0.05). For each frame, velocity fields were ignored in pixels with a low response, defined as smaller than 3-fold standard deviation (SD) of the baseline. Finally, we obtained the frequency distribution of these wave directions in each event and the average distribution of all events (see Fig. 5h).

#### Calculation of wave speed

The velocity fields calculated using the optical flow method depend on two parameters^78^ and tend to be underestimated; therefore, we used a different method to calculate the speed of waves (see Extended Data Fig. 9e). In detail, the time (T) to peak response for each pixel was determined, and the pixel with the shortest time (T_0_) to reach the peak response was defined as the origin. The wave-propagating region was then divided by fans centered at the origin with 0.5° intervals, and the relative distance (ΔS) between the distal pixel and the origin was calculated. The speed (v) in each direction was then calculated using the equation v = ΔS/(T-T_0_). Finally, we obtained the frequency distribution for speed in each event and the average distribution of all events (see Fig. 5h).

### Quantification and statistical analysis

Where appropriate, cells or animals were randomly assigned to either the control or experimental group. Imaging data were processed using ImageJ (1.53q) software (NIH) and custom-written MATLAB (R2020b) programs. Data were plotted using OriginPro 2020b (Originlab) or Adobe Illustrator CC. Except where indicated otherwise, all summary data are reported as the mean ± SEM. The SNR was calculated as the peak response divided by the SD of the baseline fluorescence fluctuation. All data were assumed to be distributed normally, and equal variances were formally tested. Differences were analyzed using the two-tailed Student’s *t*-test or one-way ANOVA; \**P*<0.05, \*\**P*<0.01, \*\*\**P*<0.001, and n.s., not significant (*P*≥0.05). Some cartoons in Fig. 3a,f, 4a, 5a and Extended Data Fig. 7a,d,g, 8a were created with BioRender.com.

## Data availability

The plasmids used to express the sensors in this study and the related sequences are available from Addgene. The human GPCR cDNA library was obtained from the hORFeome database 8.1 (http://horfdb.dfci.harvard.edu/index.php?page=home). Source data are provided with this paper.

## Code availability

The custom-written MATLAB, Arduino, and ImageJ programs will be provided upon request.

## Supporting information

5-HT and calcium waves in mouse dorsal cortex during seizure.

5-HT and endocannabinoid waves in mouse dorsal cortex during seizure.

## Acknowledgments

This work was supported by the National Key R&D Program of China (2022YFC3300905 to H.D.); the National Basic Research Program of China (2019YFE011781), the National Natural Science Foundation of China (31925017), the Beijing Municipal Science & Technology Commission (Z220009), the NIH BRAIN Initiative (1U01NS113358 and 1U01NS120824), grants from the Peking-Tsinghua Center for Life Sciences and the State Key Laboratory of Membrane Biology at Peking University School of Life Sciences, the Feng Foundation of Biomedical Research, the Clement and Xinxin Foundation, and the New Cornerstone Science Foundation (to Y.L.); the National Major Project of China Science and Technology Innovation 2030 for Brain Science and Brain-Inspired Technology (2022ZD0205600), the Postdoctoral Science Foundation (2022M720258), the Peking University Boya Postdoctoral Fellowship (to J.W.). We thank Y. Rao for sharing *Sert-Cre* mice and X. Lei at PKU-CLS and the National Center for Protein Sciences at Peking University for support and assistance with the Opera Phenix high-content screening system. We thank P. Gong at the University of Sydney and M. Mohajerani at University of Lethbridge for their help with the optical flow analysis of waves.

## Author contributions

Y.L. conceived and supervised the project. F.D., G.L., J.W. and Yu Zheng developed and optimized the sensors. F.D., J.W. and G.L. performed the experiments related to characterizing the sensors with help from X.X., Y.W., X.L. and Y.Y. J.W. performed the *in vivo* fiber photometry recordings of r5-HT sensors during optogenetic stimulation and sleep wake cycles. J.W., H.D., and L.L. performed the fiber photometry recordings of green 5-HT sensors for *in vivo* comparison during sleep wake cycles. F.D. performed the mesoscopic imaging in head-fixed mice. H.X., F.D., C.Z. and J.F. built the mesoscopic imaging system. All authors contributed to the data interpretation and analysis. F.D. and Y.L. wrote the manuscript with input from all other authors, especially the review and editing from Yulin Zhao.

---

**Extended Data Fig. 1.**
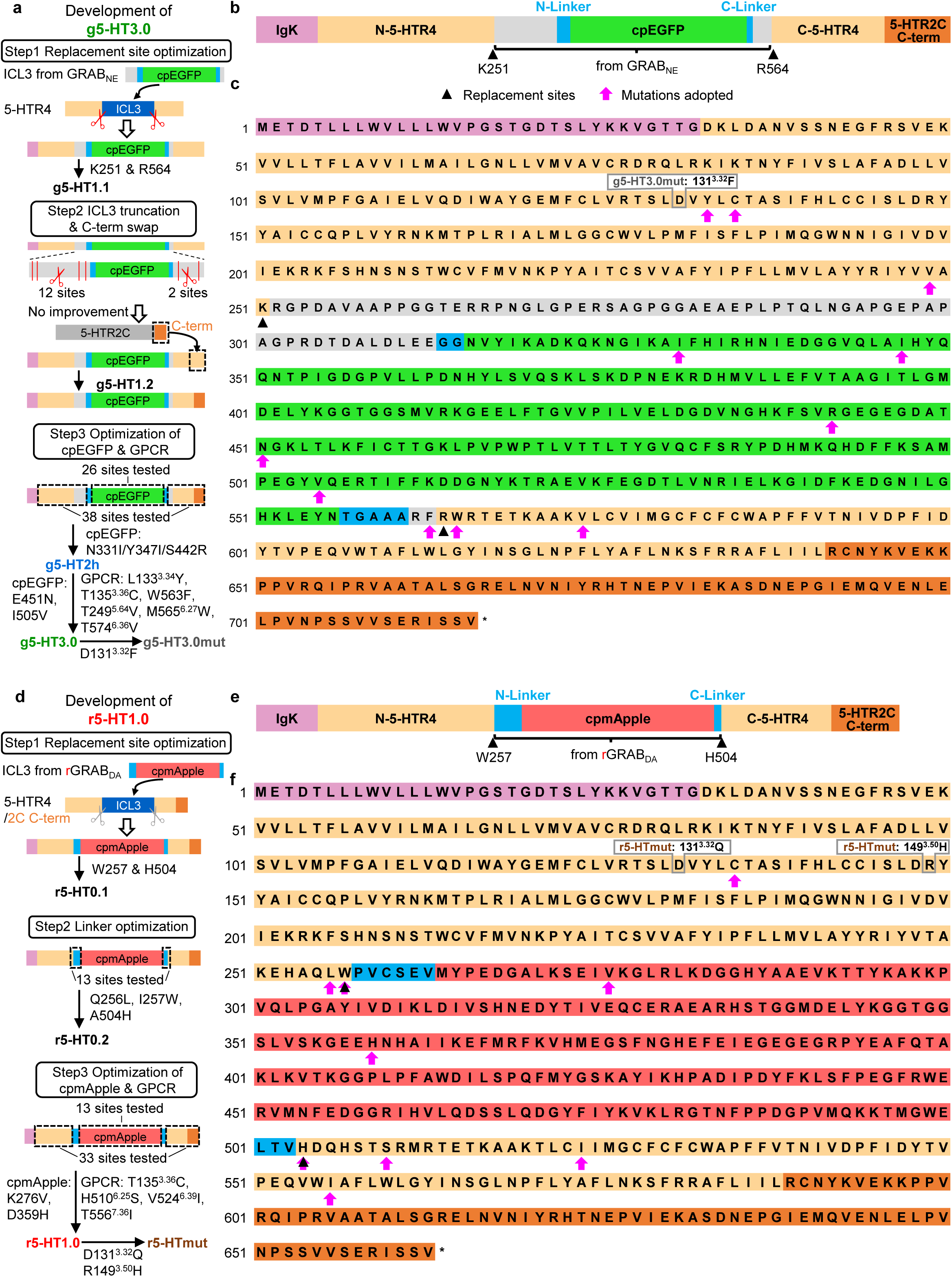
| Development and sequence of GRAB_5-HT_ sensors. **a**, A flowchart depicting the development of the g5-HT3.0 sensor, including replacement site optimization, C-terminal (C-term) swap with 5-HTR2C, linker, cpEGFP and GPCR optimization. Mutations adopted in each step are noted. **b**, Schematic showing components of the g5-HT3.0 sensor. The cpEGFP and linkers were transplanted from GRAB_NE_. **c**, The amino acid sequence of g5-HT3.0, in which replacement sites for ICL3 loop from GRAB_NE_ are denoted by black arrowhead and mutated amino acids are indicated by pink arrow. The numbering of amino acid corresponds to the start of the IgK leader in the sensor and superscripts in the insets of **a**,**c** are based on the Ballesteros-Weinstein numbering system^79^. **d–f**, The development (**d**), components (**e**) and sequence (**f**) of the r5-HT1.0 sensor. Similar to **a–c**, except that the C-term was swapped with 5-HTR2C C-term in the first step (replacement site optimization).

**Extended Data Fig. 2.**
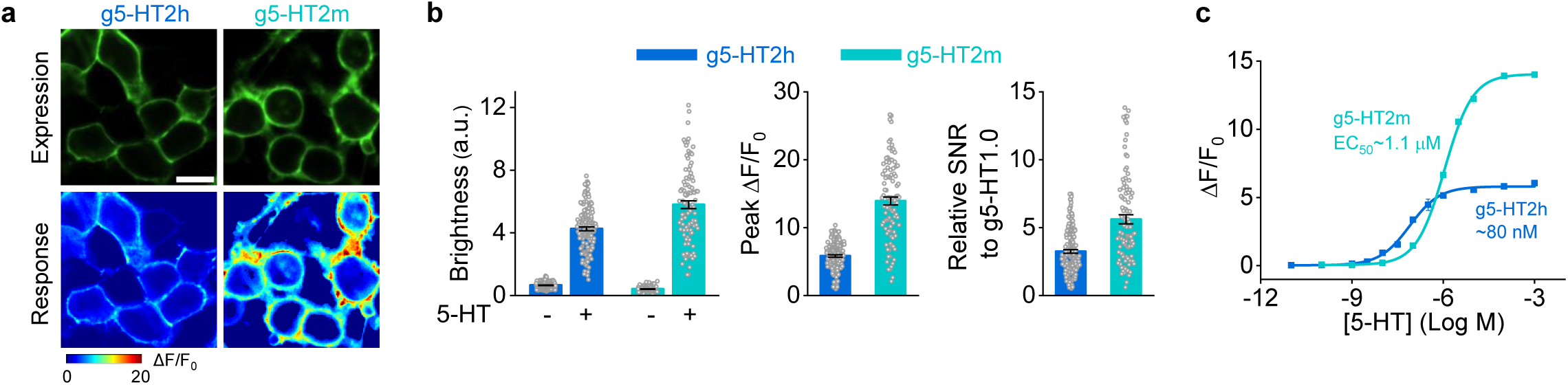
| Performance of g5-HT2h and g5-HT2m in HEK293T cells. **a**, Representative images showing the expression (top, with 5-HT) and responses (bottom) to 100 μM 5-HT for g5-HT2h (left) and g5-HT2m (right). Scale bar, 20 μm. **b**, The group summary of the brightness (left), peak ΔF/F_0_ (middle) and SNR (right) of g5-HT2h and g5-HT2m. The SNR is relative to g5-HT1.0; a.u., arbitrary units. *n* = 154 cells from 3 coverslips (154/3) for g5-HT2h, 98/3 for g5-HT2m. **c**, Dose-dependent curves of g5-HT2h and g5-HT2m. *n* = 3 wells for each sensor with 300–500 cells per well. Data are shown as mean ± SEM in **b**,**c**, with the error bars indicating the SEM.

**Extended Data Fig. 3.**
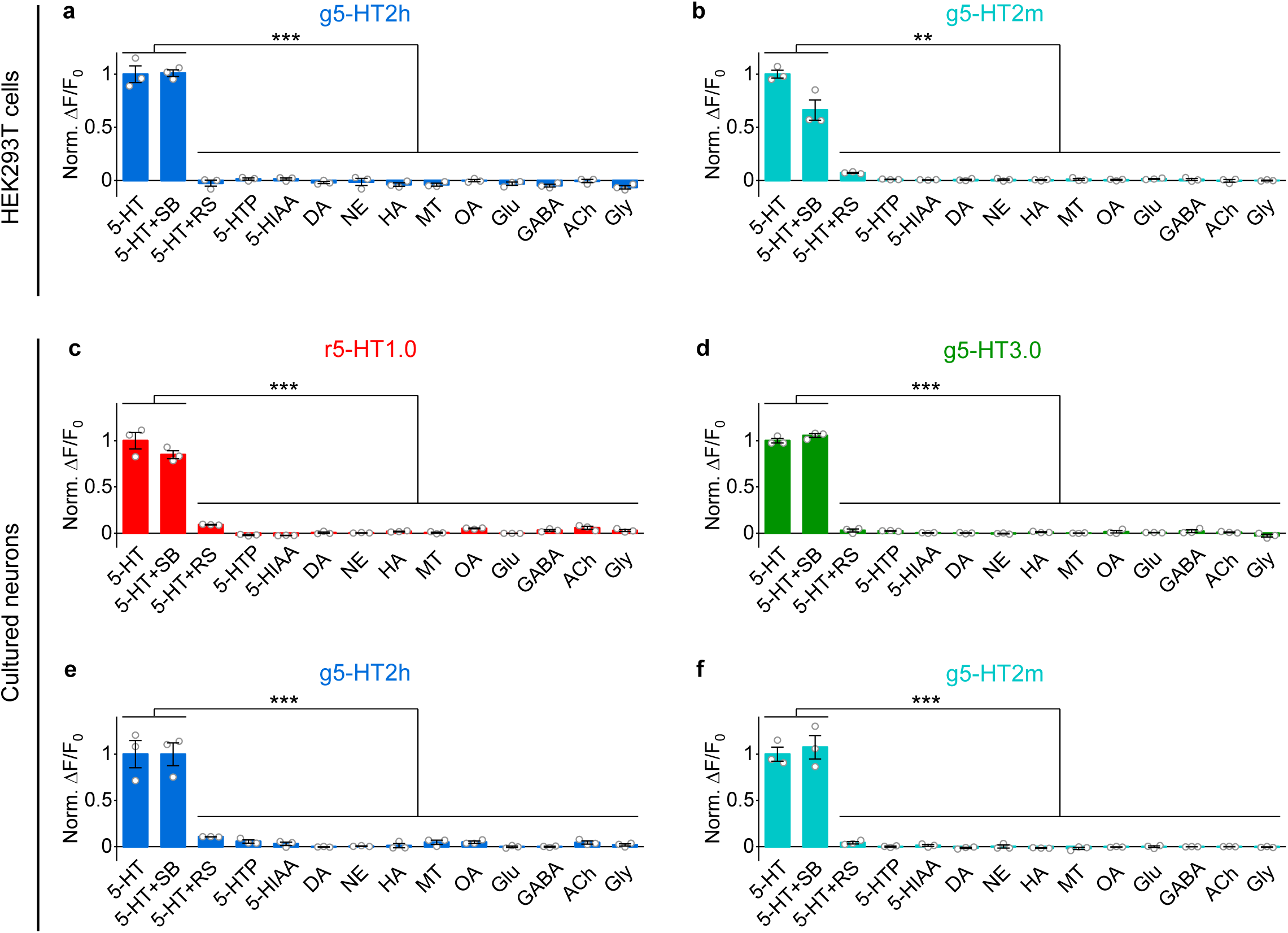
| Specificity of 5-HT sensors. Specificity test of indicated sensors in HEK293T cells (**a, b**) or cultured rat cortical neurons (**c–f**) to 5-HT alone, 5-HT together with SB, 5-HT together with RS, and 5-HT precursor, 5-HT metabolites, as well as other neurotransmitters and neuromodulators (all compounds at 10 μM except RS at 100 μM). 5-HTP, 5-hydroxytryptophan; 5-HIAA, 5-hydroxyindole acetic acid; DA, dopamine; NE, norepinephrine; HA, histamine; MT, melatonin; OA, octopamine; Glu, glutamate; GABA, gamma-aminobutyric acid; ACh, acetylcholine; Gly, glycine. Norm., normalized. *n* = 3 wells for each group with 200–500 cells per well. (One-way ANOVA followed by Tukey’s multiple-comparison tests were performed, \*\**P* < 0.01, \*\*\**P* < 0.001, n.s., not significant.) Data are shown as mean ± SEM, with the error bars indicating the SEM.

**Extended Data Fig. 4.**
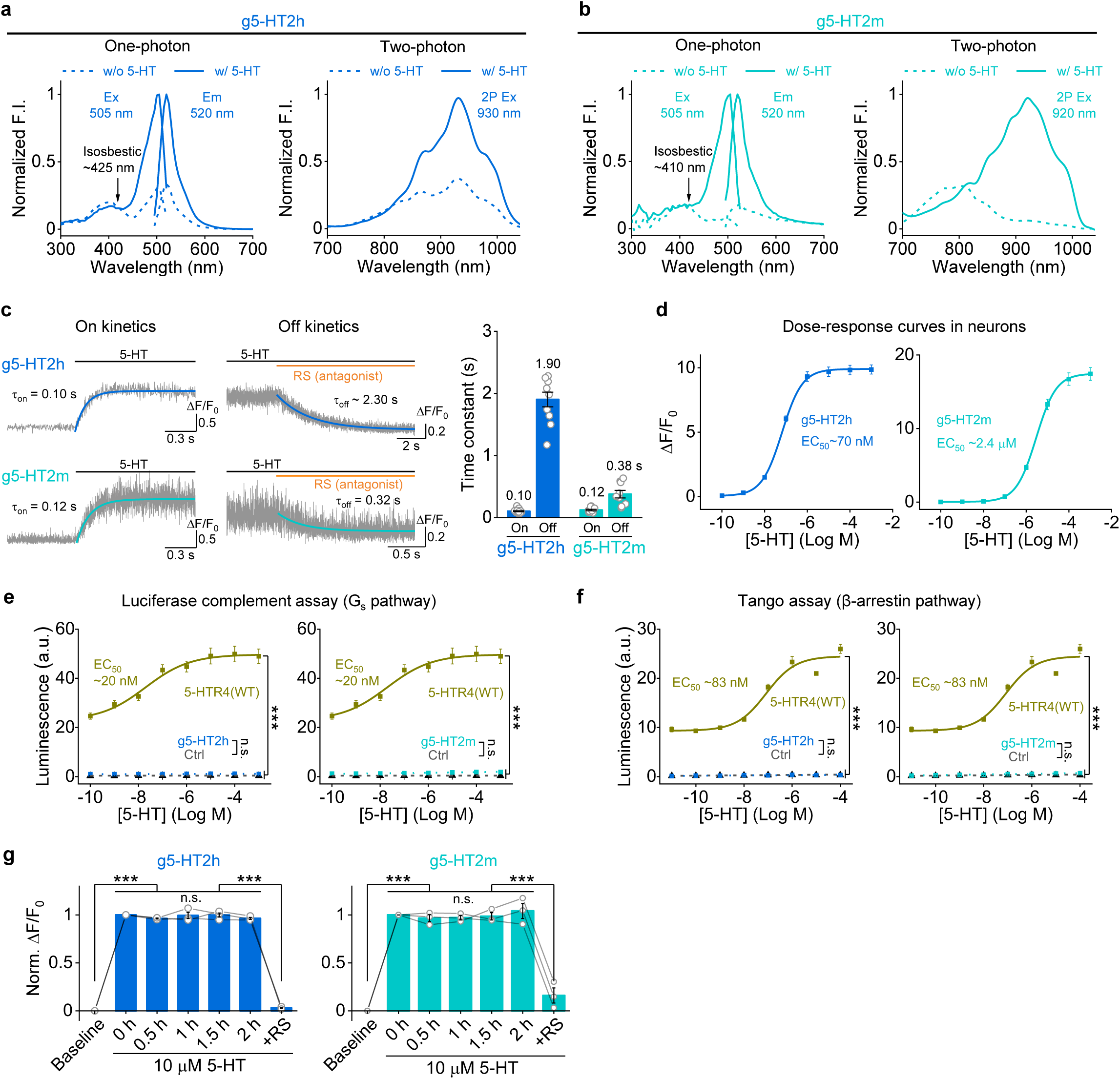
| Characterization of g5-HT2h and g5-HT2m in HEK293T cells and cultured rat cortical neurons. **a–b**, Excitation (Ex) and emission (Em) spectra of g5-HT2h (**a**) and g5-HT2m (**b**) in the absence (dash line) and presence of 10 μM 5-HT (solid line) under one-photon (left), and two-photon excitation (right). w/o, without; w/, with. **c**, Representative traces of sensor fluorescence increase to 5-HT puffing and decrease to RS puffing (left). Group summary of on and off kinetics (right). *n* = 16 cells from 4 coverslips (short for 16/4) for g5-HT2h on kinetics, 10/3 for g5-HT2h off kinetics, 11/3 for g5-HT2m on kinetics, 9/3 for g5-HT2m off kinetics. **d**, Dose-response curves of g5-HT2h (left) and g5-HT2m (right) in cultured rat cortical neurons. *n* = 60 ROIs from 3 coverslips for g5-HT2h and g5-HT2m. **e–f**, Downstream coupling tests of g5-HT2h and g5-HT2m by the luciferase complement assay for G_s_ coupling (**e**) and the Tango assay for β-arrestin coupling (**f**), respectively. WT, wild type; Ctrl, control, without expression of wild type 5-HTR4 or sensors; a.u., arbitrary units. Data of WT and Ctrl groups were replotted from Fig. 2l. *n* = 3 wells per group with 200–500 cells per well. (For luciferase complement assay, post hoc test in 1 mM 5-HT: *P* = 2.65×10^-6^ and 0.96 for g5-HT2h versus WT and Ctrl, respectively, *P* = 2.93×10^-6^ and 0.82 for g5-HT2m versus WT and Ctrl, respectively; for Tango assay, post hoc test: *P* = 4.94×10^-8^ and 1 for g5-HT2h versus WT and Ctrl, respectively, *P* = 5.96×10^-8^ and 0.88 for g5-HT2m versus WT and Ctrl, respectively.) **g**, The fluorescence of g5-HT2h (left) and g5-HT2m (right) expressed in cultured rat cortical neurons in response to a 2-h application of 5-HT, followed by 5-HTR4 antagonist RS. (For g5-HT2h, *F* = 670, *P* = 2.83×10^-5^, post hoc test: *P* = 0 for baseline versus 0 h, *P* = 0 for 2.0 h versus RS, *P* = 0.76, 1, 1, 0.80 for 0 h versus 0.5 h, 1 h, 1.5 h or 2.0 h, respectively; for 5-HT2m, *F* = 100.3, *P* = 0.006, post hoc test: *P* = 1.13×10^-6^ for baseline versus 0 h, *P* = 1.77×10^-7^ for 2.0 h versus RS, *P* = 1, 1, 1, 0.99 for 0 h versus 0.5 h, 1 h, 1.5 h or 2.0 h, respectively.) *n* = 3 wells for each sensor. One-way ANOVA (in **e**,**f**) and one-way repeated measures ANOVA (in **g**) followed by Tukey’s multiple-comparison tests, \*\*\**P* < 0.001, n.s., not significant. Data are shown as mean ± SEM in **c–g**, with the error bars indicating the SEM.

**Extended Data Fig. 5.**
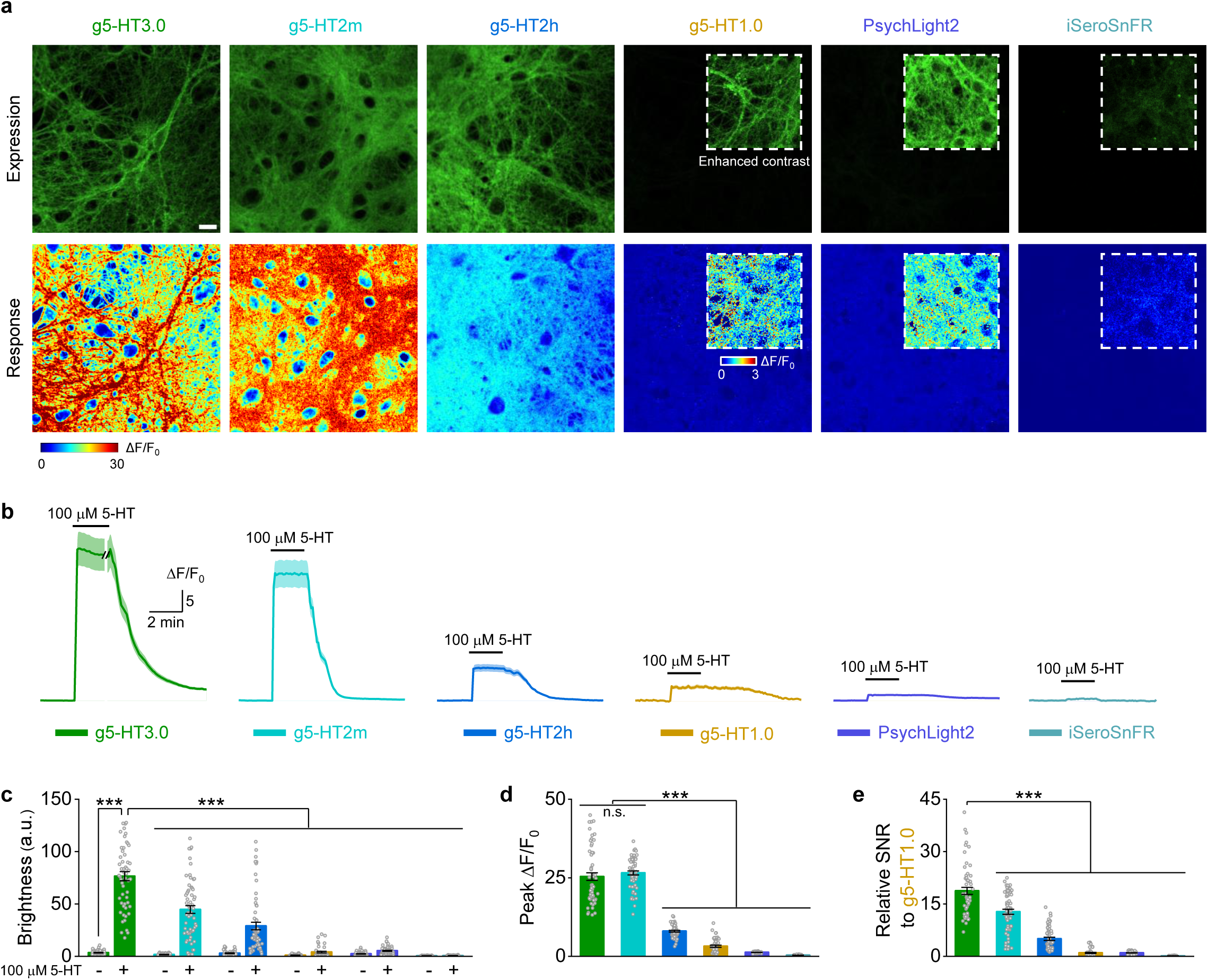
| Comparison of single GFP-based 5-HT sensors in cultured rat cortical neurons. **a**, Representative images showing the fluorescence expression (top) and responses (bottom) to 100 μM 5-HT for different sensors as indicated. Insets with white dashed outlines in images have either enhanced contrast (top) or different pseudocolor scales (bottom). Scale bar, 20 μm. **b**, Representative traces in response to 100 μM 5-HT for different sensors as indicated. **c–e**, Group summary of the brightness (**c**), peak ΔF/F_0_ (**d**) and SNR (**e**). The SNR of all sensors is relative to the SNR of g5-HT1.0; a.u., arbitrary units, the basal brightness of g5-HT1.0 is set as 1. *n* = 56 ROIs from 3 coverslip (short for 56/3) for g5-HT3.0, 60/3 for g5-HT2m, 60/3 for g5-HT2h, 48/3 for g5-HT1.0, 60/3 for PsychLight2 and 60/3 for iSeroSnFR. (One-way ANOVA followed by Tukey’s multiple-comparison tests for **c–e**; for brightness in **c**, *F*_11,676_ = 141.4, *P* = 4.97×10^-1^^67^, post hoc test: *P* <10^-7^ for g5-HT3.0 with 5-HT versus g5-HT3.0 without 5-HT and other sensors with or without 5-HT; for peak ΔF/F_0_ in **d**, *F*_5,338_ = 446.9, *P* = 1.46×10^-1^^46^, post hoc test: *P* =0.696 for g5-HT3.0 versus g5-HT2m, *P* <10^-7^ for g5-HT3.0 and g5-HT2m versus other sensors; for relative SNR in **e**, *F*_5,338_ = 195.1, *P* = 2.46×10^-97^, post hoc test: *P* <10^-7^ for g5-HT3.0 versus other sensors, \*\*\**P* < 0.001, n.s., not significant.) Data are shown as mean ± SEM in **b–e**, with the error bars or shaded regions indicating the SEM.

**Extended Data Fig. 6.**
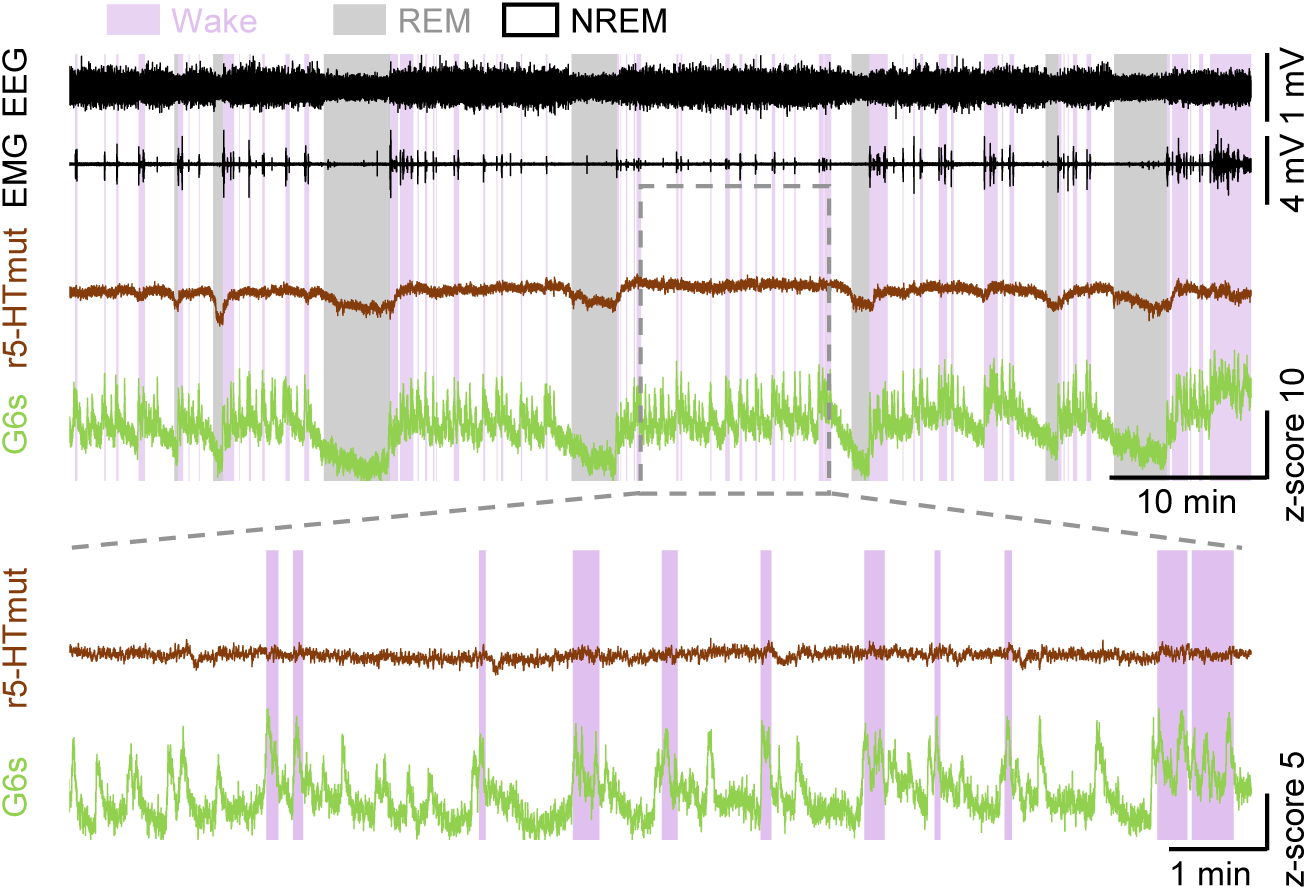
| Representative r5-HTmut and GCaMP6s signals during the sleep-wake cycle in freely moving mice. Representative r5-HTmut and GCaMP6s (G6s) traces in the mouse basal forebrain (BF) along with EEG and EMG recording during the spontaneous sleep-wake cycle.

**Extended Data Fig. 7.**
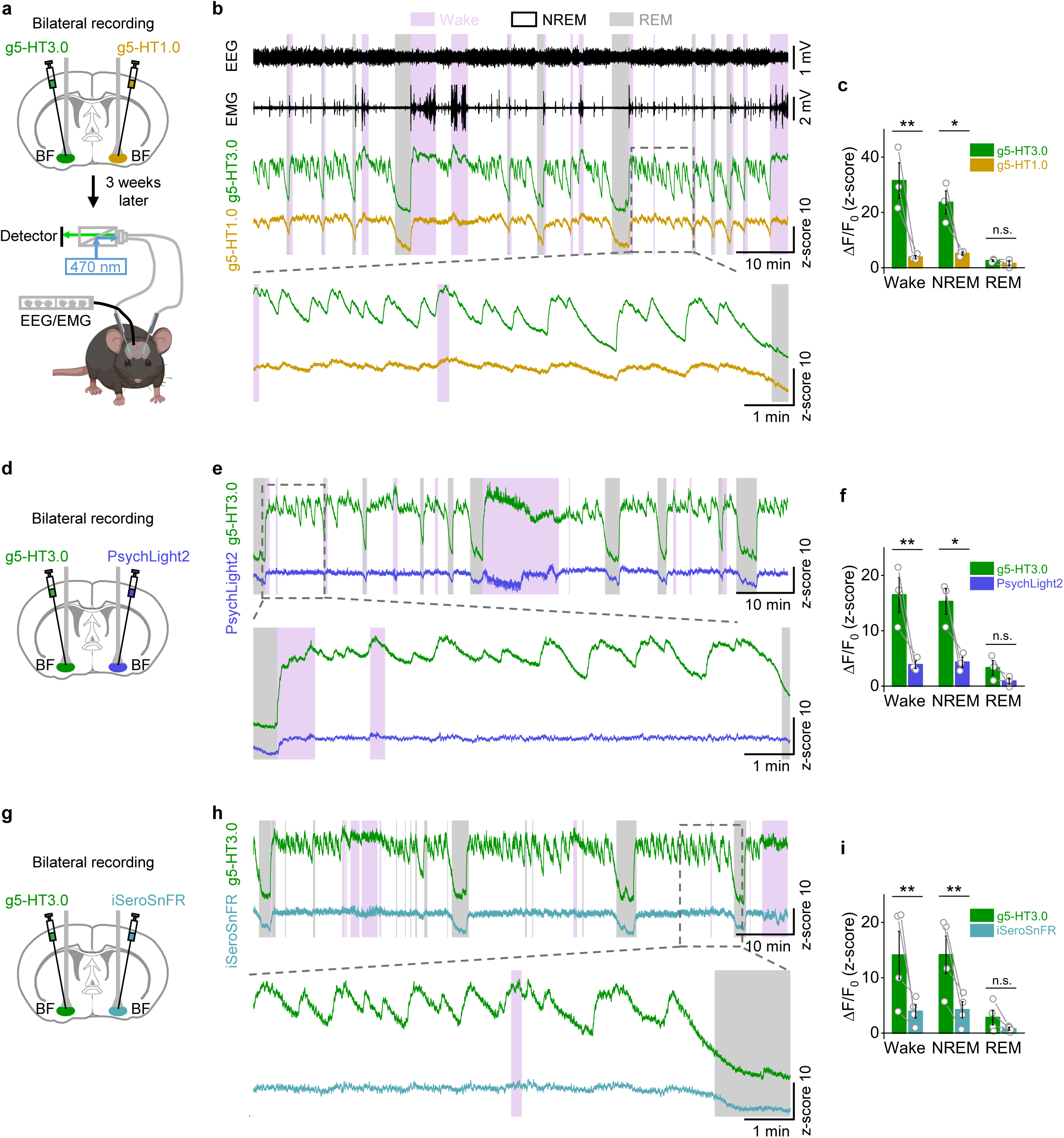
| Comparison of gGRAB_5-HT3.0_ and other green 5-HT sensors during the sleep-wake cycle in freely moving mice. **a**, Schematic showing the setup of bilateral fiber-photometry recording of g5-HT3.0 and g5-HT1.0 during the sleep-wake cycle in mice. **b**, Representative traces of simultaneous EEG, EMG, g5-HT3.0 and g5-HT1.0 recording during the sleep-wake cycle in freely behaving mice. Pink shading, wake state; gray shading, REM sleep. **c**, Summary of averaged g5-HT3.0 and g5-HT1.0 signals in indicated sleep-wake states. *n* = 3 mice. (*P* = 0.0034, 0.014 and 0.83 during wake, NREM and REM sleep state, respectively.) **d–f**, Similar to **a–c**, except bilateral recording of g5-HT3.0 and PsychLight2, *n* = 3 mice in **f**. (*P* = 0.0066, 0.011 and 0.38 during wake, NREM and REM sleep state, respectively.) **g–i**, Similar to **a–c**, except bilateral recording of g5-HT3.0 and iSeroSnFR, *n* = 4 mice in **i**. (*P* = 0.0086, 0.0095 and 0.47 during wake, NREM and REM sleep state, respectively.) Data are shown as mean ± SEM in **c**,**f**,**i**, with the error bars or shaded regions indicating the SEM. Two-way repeated measures ANOVA followed by Tukey’s multiple-comparison tests in **c**,**f**,**i**, \**P* < 0.05, \*\**P* < 0.01, n.s., not significant.

**Extended Data Fig. 8.**
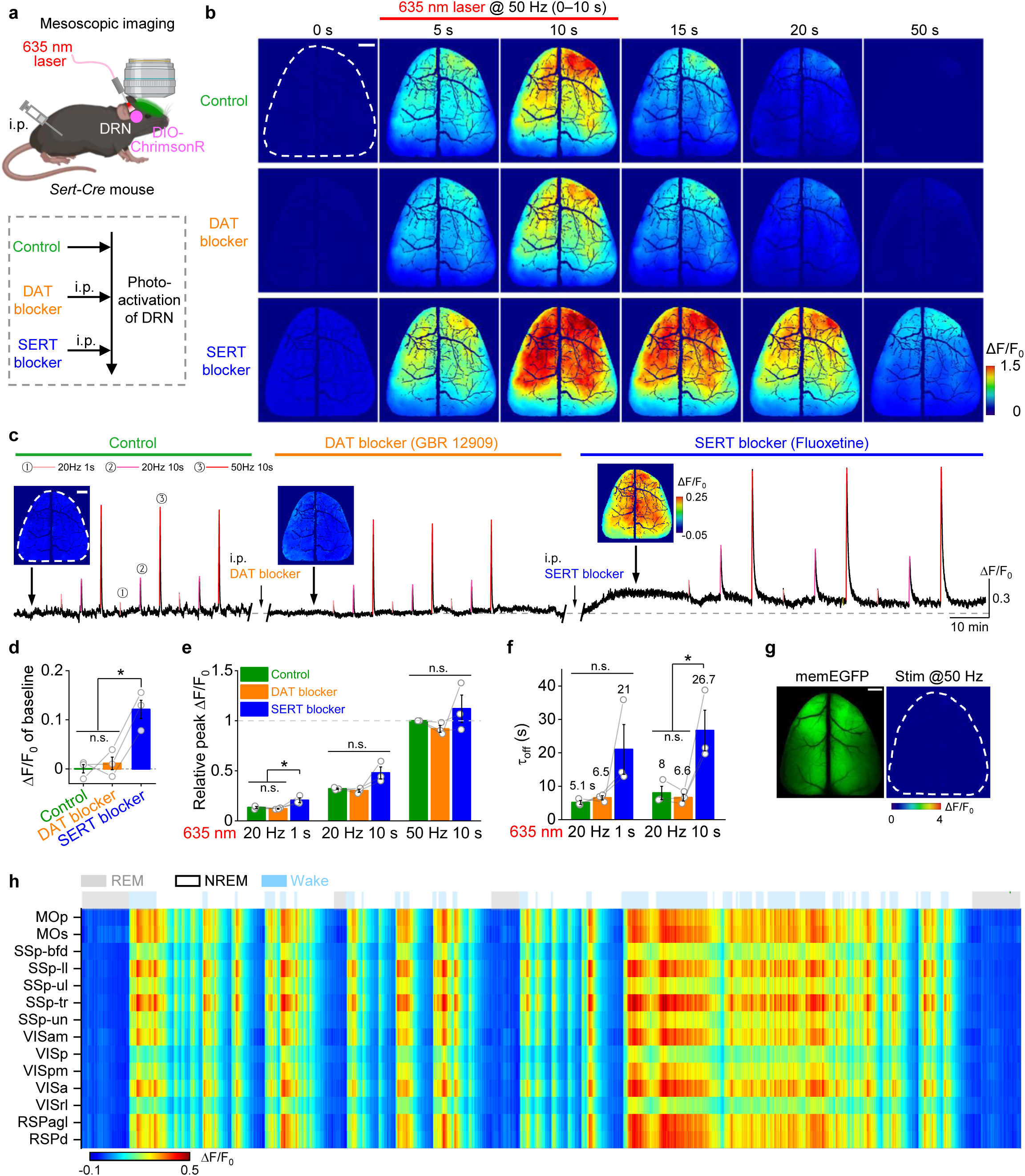
| gGRAB_5-HT3.0_ reveals 5-HT dynamics in mouse dorsal cortex *in vivo*. **a**, Schematic depicting the protocol for mesoscopic imaging along with optogenetic activation of DRN with different drug treatments. **b**, Representative pseudocolor images in response to the 50 Hz 10 s optical stimulation of DRN with indicated treatments. **c**, Representative trace of g5-HT3.0 with indicated treatments, including the application of different drugs and activation of DRN using a 635 nm laser with different frequencies and durations. Insets above the trace are averaged images in the indicated baseline timepoint (by the black arrow) of different stages. **d**, Group data of averaged g5-HT3.0 baseline fluorescence changes under indicated treatments. (One-way repeated measures ANOVA followed by Tukey’s multiple-comparison tests, *F* = 19.9, *P* = 0.047, post hoc test: *P* = 0.896 for control versus DAT blocker, 0.016 for SERT blocker versus control and 0.022 for SERT blocker versus DAT blocker.) **e–f**, Group summary of optical stimulation evoked peak response (**e**) and decay kinetics (**f**). *n* = 3 mice in **d–f**. (One-way repeated measures ANOVA followed by Tukey’s multiple-comparison tests. For relative peak ΔF/F_0_ in **e**, under 20 Hz 1 s stimulation, *F* = 11.1, *P* = 0.023, post hoc test: *P* = 0.81 for control versus DAT blocker, 0.043 for SERT blocker versus control and 0.026 for SERT blocker versus DAT blocker; under 20 Hz 10 s stimulation, *F* = 6.67, *P* = 0.053; under 50 Hz 10 s stimulation, *F* = 1.39, *P* = 0.348. For decay kinetics ԏ_off_ in **f**, under 20 Hz 1 s stimulation, *F* = 4.06, *P* = 0.182; under 20 Hz 10 s stimulation, *F* = 16.78, *P* = 0.011, post hoc test: *P* = 0.932 for control versus DAT blocker, 0.018 for SERT blocker versus control and 0.014 for SERT blocker versus DAT blocker.) **g**, Representative images showing the memEGFP expression and response to the 50 Hz 10s optical activation. **h**, Representative heatmap showing changes of g5-HT3.0 fluorescence in different brain regions during the sleep-wake cycle. Gray shading, REM sleep; light blue shading, wake state. The dashed white outlines **b**,**c**,**g** indicate the ROI. All scale bar, 1 mm. Data are shown as mean ± SEM in **d–f**, with the error bars indicating the SEM, \**P* < 0.05, n.s., not significant.

**Extended Data Fig. 9.**
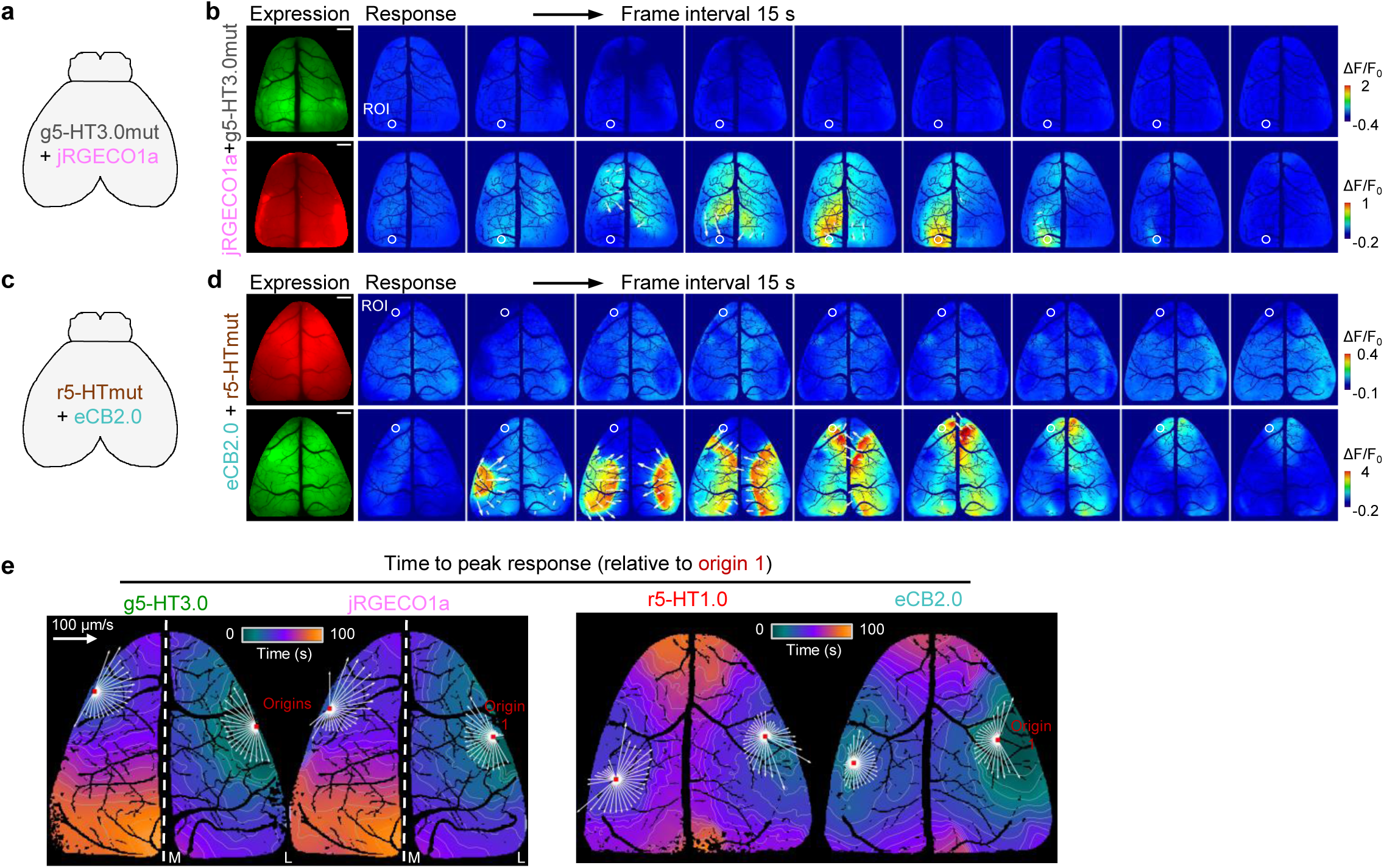
| Mesoscopic imaging of 5-HT, Ca^2+^ and eCB waves during seizures. **a**, Schematic showing the co-expression of g5-HT3.0mut and jRGECO1a in the mouse dorsal cortex. **b**, Representative images show fluorescence changes of g5-HT3.0mut (top) and jRGECO1a (bottom) during seizures. A ROI labeled with the white circle (500 μm in diameter) shows the maximum response regions of jRGECO1a, which corresponds to the trace in Fig. 5c. White arrows indicate the direction of wave propagation and the length of arrows indicates relative magnitudes of velocities. Scale bar, 1 mm. **c–d**, Similar to **a–b**, but co-expressing r5-HTmut and eCB2.0. The ROI shows the maximum response regions of eCB2.0 and corresponds to the trace in Fig. 5e. **e**, Representative time to peak response maps of waves relative to the origin 1, monitored by different sensors. Red dots indicate origin locations of waves; white arrows indicate velocity vectors calculated based on the propagation distance and duration along the corresponding direction; L, lateral, M, medial; scale bar of speed, 100 µm/s.

